# Immunoinformatics design of a novel multiepitope vaccine candidate against non-typhoidal salmonellosis caused by *Salmonella* Kentucky using outer membrane proteins A, C, and F

**DOI:** 10.1101/2024.06.03.597183

**Authors:** Elayoni E. Igomu, Paul H. Mamman, Jibril Adamu, Maryam Muhammad, Abubarkar O. Woziri, Manasa Y. Sugun, John A. Benshak, Kingsley C. Anyika, Rhoda Sam-Gyang, David O. Ehizibolo

## Abstract

The global public health risk posed by *Salmonella* Kentucky, particularly due to the dissemination of antimicrobial resistance genes in human and animal populations, is rising. This serovar, widespread in Africa, has emerged as a notable cause of non-typhoidal gastroenteritis in humans. In this study, we utilized a bioinformatics approach to design a peptide-based vaccine targeting epitopes from the outer membrane proteins A, C, and F of *Salmonella* Kentucky, with the *fliC* protein from *Salmonella* Typhimurium serving as an adjuvant. Through this approach, we identified 14 CD8+ and 7 CD4+ T-cell epitopes, which are predicted to be restricted by various MHC class I and MHC class II alleles. When used in vaccine formulations, the predicted epitopes are expected to achieve a population coverage of 94.91%. Additionally, we identified seven highly immunogenic linear B-cell epitopes and three conformational B-cell epitopes. These T-cell and B-cell epitopes were then linked using appropriate linkers to create a multi-epitope vaccine (MEV). To enhance the immunogenicity of the peptide construct, the *fliC* protein from *Salmonella* Typhimurium was included at the N-terminal as an adjuvant. The resulting MEV construct demonstrated high structural quality and favorable physicochemical properties. Molecular docking studies with Toll-like receptors 1, 2, 4, and 5, followed by molecular dynamic simulations, suggested that the vaccine-receptor complexes are energetically feasible, stable, and robust. Immune simulation results showed that the MEV elicited significant responses, including IgG, IgM, CD8+ T-cells, CD4+ T-cells, and various cytokines (IFN-γ, TGF-β, IL-2, IL-10, and IL-12), along with a noticeable reduction in antigen levels. Despite these promising in silico results, further validation through preclinical and clinical trials is necessary to confirm the vaccine’s protective efficacy and safety.

## 1.0 Introduction

The role of antibiotics in modern medicine is indispensable. They are crucial not only for treating various infectious diseases but also as an essential component of supportive therapy in cancer treatments. Additionally, antibiotics are vital following surgical procedures and play a significant role in livestock and food animal production [1]. However, the widespread and improper use of antibiotics has significantly contributed to the rise of antibiotic resistance in pathogenic bacteria, bringing us closer to the grim reality of a post-antibiotic era. Predictions suggest that by 2050, deaths caused by antimicrobial-resistant bacteria could surpass those caused by cancer [1].

According to the World Health Organization’s (WHO) bacterial priority pathogen list, fluoroquinolone-resistant non-typhoidal *Salmonella*, alongside third-generation cephalosporin-resistant and carbapenem-resistant *Enterobacter* species, are among the highest priority pathogens posing significant risks to public health [2]. Notably, All of these high-level resistances have been detected in *Salmonella enterica* subsp. *enterica* serovar Kentucky (*S.* Kentucky), together with well-documented zoonotic potential, and instances of *S.* Kentucky being reported across various animal species and humans, makes *S.* Kentucky a high-risk global Multidrug-resistant (MDR) pathogen [3, 4]. The emergence, rising cases of reports, and continual spread of MDR *S.* Kentucky has been highlighted by different health organizations and nations and its continuous association with high-level fluoroquinolone resistance has gained epidemiological significance globally [5]. For severe non-typhoidal *Salmonella* infections in both humans and animals, the preferred treatments are ciprofloxacin and third-generation cephalosporins. However, the resistance of *S.* Kentucky to these frontline antibiotics has greatly narrowed the available treatment options [3].

Vaccines have been instrumental for decades in the prevention of diseases and have had an unprecedented impact on human and animal health globally. Unlike antibiotics, vaccines have shown a much lower propensity for contributing to antimicrobial resistance (AMR) [7]. This difference arises because vaccines and antibiotics work in fundamentally different ways. Vaccines are primarily used as a preventive measure, taking action before the invading bacterial pathogen can multiply. This preemptive approach reduces both the pathogen load and the extent of tissue invasion, substantially lowering the chances of AMR developing or antimicrobial-resistant mutations occurring [8].

*Salmonella* outer membrane proteins (OMPs) have been investigated as potential vaccine candidates over the past several years. The *Salmonella* OMPs viz outer membrane protein A (OmpA), outer membrane protein C (OmpC), and outer membrane protein F (OmpF), are widely distributed and conserved across various *Salmonella* serovars. This conservation suggests their potential use in diagnostics or as components of subunit or conjugate vaccines [9, 10]. Several studies have used crude preparation of *Salmonella* OMPs to evoke a strong immunological response, thereby conferring protective immunity against an array of Gram-negative bacteria [10]. Notably, OMPs possess surface-exposed epitopes that are easily recognized by T-cell and B-cell receptors. [11, 12].

Recently, the WHO underscored the urgency of developing new vaccines against infectious bacterial pathogens in Sub-Saharan Africa [13, 14], with a specific focus on non-typhoidal salmonellosis (NTS). This region is particularly vulnerable to the emergence of antimicrobial resistance due to various risk factors [3, 15, 16, 17]. In response to this call, our study aims to design a multiepitope vaccine candidate against NTS caused by *Salmonella* Kentucky. Using computational immunological methods, we retrieved sequences of OmpA, OmpC, and OmpF from the National Center for Biotechnology Information (NCBI) databank to inform our vaccine design

## 2.0 Materials and Methods

### 2.1 Retrieval of outer membrane proteins (OMPs) of *Salmonella* Kentucky sequences

FASTA format sequences for OmpA, OmpC, and OmpF of *S*. Kentucky strains were retrieved from the NCBI database (https://www.ncbi.nlm.nih.gov/protein/). All the accession numbers of OMPs of *S*. Kentucky used were noted. To maximize the similarity between proteins of the different strains of *S*. Kentucky OMPs, the retrieved sequences were blasted with a query set at 99 % to 100 % identity (BLAST: https://blast.ncbi.nlm.nih.gov). A FASTA sequence of flagellin protein (*fliC*) (GenBank accession number KAA0422324.1) of *Salmonella enterica subsp. enterica* serovar Typhimurium (*Salmonella* Typhimurium) was chosen as an adjuvant to enhance the efficacy of the vaccine peptide [18, 19].

### 2.2 Multiple sequence alignment of OMPs and generation of consensus sequence

Generating a consensus sequence for each class of OMPs was paramount to ensure protein residue conservation. The numerous OMPs FASTA sequences retrieved were aligned using Jalview (2.11.2.7). Jalview software is a free cross-platform program for multiple sequence alignment editing, visualization, and analysis (https://www.jalview.org/ [20]. The FASTA sequences of all selected Omp A, C, and F from the NCBI database were manually entered into the Jalview software program which automatically aligns and generates predicted consensus sequences using Blosum62 parameters [21].

### 2.3 Epitope mapping

To determine what major histocompatibility complex (MHC) epitopes are to be used for the vaccine design. Each of the consensus sequences was subjected to epitope mapping prediction on the Immune Epitope Database (IEDB) server (http://tools.iedb.org/main/tcell). The IEDB server contains MHC binding data related to major histocompatibility complex class I (MHC-I) and major histocompatibility complex class II (MHC-II) [22, 23].

#### 2.3.1 Cytotoxic T-lymphocytes (CTL) epitopes prediction

Cytotoxic T-lymphocytes also called CD8+ T-cells mostly express T-cell receptors (TCRs) capable of recognizing distinctive antigens. Thus, Identifying cytotoxic T-lymphocyte epitopes forms an integral part of designing a multiepitope vaccine. The prediction of cytotoxic T-lymphocytes activating epitopes that bind to MHC-I was predicted on the IEDB recommended 2023.09 (NetMHCpan4.1 EL) server (http://tools.iedb.org/mhci/). This server utilizes the weight matrix of the MHC-I binding peptides, the transporter associated with antigen processing (TAP), and the proteasomal C-terminus cleavage score for its prediction. The MHC source species was set as Human and the HLA allele reference set was predicted with 9-mer epitope length. The threshold for the classification of epitopes as strong or weak binders was set at 0.5 % and 0.2 % respectively [23, 24].

#### 2.3.2 Helper T-lymphocytes (HTL) epitopes prediction

One the critical components of the cellular and humoral immune system are helper T-lymphocytes also called CD4+ T-cells. They are key regulators of a cascade of inflammatory processes necessary to subdue infection [25] and therefore form an integral part of multiepitope vaccine design. The prediction of helper T-lymphocytes activating epitopes that bind to MHC-II was predicted on the IEDB recommended 2023.09 (NetMHCpan4.1 EL) server (http://tools.iedb.org/mhcii/). This server provides a ranked list of peptides based on calculations of IC50 values for the predicted epitopes, which are inversely related to their binding affinity to MHC-II molecules. IC50 values below 50 nM indicate high binding affinity, values below 500 nM suggest intermediate binding affinity and values below 5000 nM indicate low binding affinity of the epitopes to MHC-II. The 7-allele human leukocyte antigen (HLA) reference set was utilized to predict epitopes, with HLA-DR selected specifically for Homo sapiens. Subsequently, a 15-mer length epitope with a percentile rank classification was obtained using the IEDB recommended 2023.05 method (NetMHCIIpan 4.1L) [18, 22, 24].

### 2.4 Prediction of linear B-lymphocyte epitopes

The initiation of humoral immune responses requires antigen-reactive B-lymphocytes (B-cells) to encounter an antigen. B-lymphocytes, unlike T-lymphocytes, have B-cell receptors (BCRs), which bind to foreign antigens and trigger an antibody response. These BCRs are highly specific, with all BCRs on a single B-cell recognizing the same epitope. B-lymphocyte epitopes were predicted using the ABCpred server (https://webs.iiitd.edu.in/raghava/abcpred/ABC_submission.html) and a neural network algorithm. A sequence length of 16 amino acids and a binding score threshold of > 0.51 were employed to predict linear B-cell epitopes for the OmpA, OmpC, and OmpF proteins using their FASTA consensus sequences. Epitopes with higher scores were selected for vaccine construction. Additionally, the Linear B-cell epitopes were further assessed for antigenicity using the antigen sequence properties tool on the IEDB website (http://tools.iedb.org/bcell/result/) [18, 19].

### 2.5 Prediction of antigenicity, allergenicity and toxicity

The antigenicity of each predicted CTL, HTL, and linear B-cell epitope, as well as the constructed multi-epitope vaccine sequence, was assessed using the VaxiJen v2.0 server (http://www.ddg-pharmfac.net/vaxijen/VaxiJen/VaxiJen.html), with a threshold set at 0.4 for bacteria. This server employs auto and cross-covariance (ACC) transformation to generate its output [26, 27, 28]. To evaluate the potential allergenicity of the determined epitopes, the AllerTOP v2.0 server (http://www.ddg-pharmfac.net/AllerTOP) was utilized. AllerTOP also uses ACC transformation to convert protein sequences into uniform, equal-length vectors. It then compares the E descriptors of amino acids to obtain the closest k value. These k values are obtained based on a training set of 2,210 known allergens from various species and 2,210 non-allergens from the same species [29, 30]. The predicted CTL, HTL, and linear B-cell epitopes was subjected to analysis on the ToxinPred server (https://webs.iiitd.edu.in/raghava/toxinpred/multi_submit.php), with parameters set at default to evaluate their toxicity. The ToxinPred server (https://webs.iiitd.edu.in/raghava/toxinpred/multi_submit.php) uses several algorithms to predict peptide toxicity. These include Quantitative Matrix (QM) methods, motif-based methods, and machine learning models such as Support Vector Machines (SVM), Random Forests (RF), and Decision Trees (DT). These approaches analyze peptide sequences to assess their potential toxicity [23, 27].

### 2.6 Interferon-gamma (IFN-γ) inducing epitope prediction

Interferon-gamma (IFN-γ) is critical for pathogen recognition and elimination, acting as the central effector of cell-mediated immunity [31].This cytokine induction is a vital consideration for designing a multiepitope vaccine (MEV) peptide against an infectious disease. The presence of IFN-γ inducing epitopes in the proposed MEV peptide was predicted using the IFNepitope server (http://crdd.osdd.net/raghava/ifnepitope/index.php). The server, trained with MHC-II epitopes, generates all possible overlapping peptides from an antigen based on IFN-γ inducing and non-inducing datasets, employing a Support Vector Machine (SVM) prediction system. The 15-mer MHC-II epitopes were submitted to the IFN-γ epitope server to determine if the HTL epitopes could induce an IFN-γ immune response. A hybrid motif and SVM model were selected from the available algorithm models, with a prediction score threshold set at >1.0 for epitope selection. Ultimately, high-response IFN-γ epitopes were selected for the MEV [32, 33].

### 2.7 Epitope conservancy analysis

Epitope conservancy is the degree to which a particular epitope sequence is preserved across different strains or species of a pathogen. The conservation of epitopes was predicted using the IEDB Conservancy Analysis Tool (http://tools.iedb.org/conservancy/), a web-based tool designed to analyze epitope conservancy across various protein sequences [34]. Predicted CTL and HTL epitope sequences, along with their corresponding OMP class sequences, were entered into their respective text areas and the minimum conservancy sequence identity threshold was set at ≥ 100% [35].

### 2.8 Population coverage analysis of MHC-I and MHC-II epitopes

The IEDB’s Population Coverage tool (http://tools.iedb.org/population/) was used to analyze how the affinity of CTL and HTL epitopes for HLA alleles varies by race, location, and country and how it impacts our multiepitope peptide vaccine design. The IEDB Population Coverage tool estimates the proportion or distribution of individuals expected to exhibit a response to the specified epitopes according to their identified HLA profiles. Furthermore, it evaluates the mean number of epitope matches/HLA allele pairings acknowledged by the overall populace, alongside the maximum and minimum counts of epitope matches acknowledged by 90% of the chosen population. Using the default parameters, 16 geographical areas were selected for the population coverage analysis [36, 37].

### 2.9 Construction of multiepitope vaccine sequence

For the final assembly of the multi-epitope vaccine, epitopes with high antigenicity, absence of allergenicity, and non-toxic properties were selected. B-cell epitopes (OmpA, OmpC, and OmpF) and HTL epitopes were connected using a GPGPG linker, while CTL epitopes were linked with an AAY linker. To further augment the immunogenicity of the multi-epitope vaccine, the adjuvant flagellin (*fliC)* protein (GenBank accession number KAA0422324.1) from *Salmonella enterica subsp. enterica* serovar Typhimurium was incorporated via the EAAAK linker at the N-terminal of the epitopes [18, 38, 39].

### 2.10 Prediction of physicochemical characteristics and solubility

The MEV structure underwent analysis on the Expasy Protparam online server (https://web.expasy.org/protparam/) to forecast various physicochemical traits of the multi-peptide arrangement, including the aliphatic index, molecular weight (MW), theoretical isoelectric point (pI), half-life, instability, and grand average of hydropathicity (GRAVY) [40, 41]. Additionally, the solubility value of the MEV was determined using the Protein-Sol server (http://protein-sol.manchester.ac.uk). A scaled solubility value exceeding 0.45 indicates a higher solubility profile compared to the average soluble *Escherichia coli* (*E. coli*) protein in the experimental solubility dataset. Conversely, proteins with lower scaled solubility values are expected to be insoluble [42].

### 2.11 Secondary structure prediction

The secondary structure of a protein consists of locally folded formations within a polypeptide, driven by interactions among backbone atoms. These formations include α β -sheets), and random coils. To determine the secondary structure of the MEV construct, predictions were made using the online tools PSIPRED and SOPMA. PSIPRED (http://bioinf.cs.ucl.ac.uk/psipred/) employs a neural network-based approach to assess the -helix, β strand, or coil [43]. SOPMA (https://npsa.lyon.inserm.fr/cgi-bin/npsa_automat.pl?page=/NPSA/npsa_sopma.html) similarly predicts secondary structures. Utilizing both PSIPRED and SOPMA allows for a more comprehensive and accurate prediction of a protein’s secondary structure by capitalizing on the strengths of each tool [44].

### 2.12 Tertiary structure prediction

To precisely forecast and model the three-dimensional (3D) configuration of the vaccine peptide and analyze the protein sequences’ domains, the I-TASSER (Iterative Threading Assembly Refinement) software (https://zhanglab.ccmb.med.umich.edu/I-TASSER) was employed. This software is well-regarded for its accuracy in predicting 3D protein structures. It utilizes a sophisticated approach involving multiple threading alignments to identify template structures from the Protein Data Bank (PDB). Subsequently, it assembles full-length models by integrating structural fragments from these templates. Further refinement of the predicted models is achieved through iterative simulations to optimize their structure. The confidence score (C-score) is utilized to assess the accuracy of the 3D model, with higher values indicating superior quality, with the general range between −5 and 2 [19, 33, 45].

### 2.13 Refinement and validation of tertiary structure

Validating the tertiary structure is essential in vaccine development to identify potential flaws in the predicted model [46]. The GalaxyRefine web server (http://galaxy.seoklab.org/cgi-bin/submit.cgi?type=REFINE) was used to enhance the quality of the 3D structure. This tool uses a sophisticated routine that involves reassembly and molecular dynamics modulation to optimize local structural regions, particularly improving side-chain conformations and main-chain positioning. The iterative refinement process of the server utilizes a force field that balances attractive and repulsive forces, resulting in a more stable and realistic structure. The server produces several refined models, ranked by quality, and with metrics such as RMSD, MolProbity scores, and other structural indicators provided for assessment [47]. The ProSA-web server was employed to assess the overall quality score of the precise input structure for 3D validation. Additionally, ERRAT (http://services.mbi.ucla.edu/ERRAT/) was utilized to analyze unbonded inter-atomic interactions and high-resolution crystallographic structures [48, 49]. To validate the refined tertiary structure of the vaccine model, the PROCHECK server (https://saves.mbi.ucla.edu) and VADAR version 1.8 (http://vadar.wishartlab.com/index.html?) were used [50]. The PROCHECK server validates protein structure quality by evaluating dihedral angles, bond lengths, bond angles, and geometric properties. It provides detailed stereo-chemical statistics and comprehensive reports, ensuring the accuracy and reliability of protein models. The outcome from these servers includes the Ramachandran plot, illustrating the proportion of residues situated within favored, permitted, and disallowed regions [51].

### 2.14 Prediction of discontinuous B-cell epitopes

The folding of proteins can bring distant residues into proximity, leading to the formation of discontinuous B-cell epitopes. The prediction of these discontinuous epitopes relies on the Protrusion Index (PI) value and the clustering based on the distance R, which measures the distance between the center of mass of the residue. A higher R-value indicates a higher likelihood of encountering discontinuous epitopes. Given that more than 90% of B-cell epitopes are discontinuous, the Ellipro server (http://tools.iedb.org/ellipro/) was utilized to predict the presence of these epitopes within the vaccine structure [23, 52]. Default threshold parameters were applied, with a Minimum residue score of 0.5 and a Maximum distance of 6 A. This server evaluates the 3D structure of the vaccine and assigns an ellipsoid score to each residue, represented by the PI value. A PI score of 0.9 implies that 90% of the protein’s residues fall within the ellipsoid, while the remaining 10% are outside of it. The determination of this PI value considers the center of mass of each residue, which is situated outside of the largest feasible ellipsoid [23, 28].

### 2.15 Protein-protein docking

To elicit an immune response, the vaccine model is expected to bind with immunological receptors. Toll-like receptors (TLRs) such as TLR-1, TLR-2, TLR-4, and TLR-5 play crucial roles in recognizing *Salmonella* structures and triggering the production of inflammatory cytokines due to their high sensitivity to bacterial components like triacylated lipoproteins (TLR-1), lipoproteins (TLR-2), lipopolysaccharide (TLR-4), and flagellin (TLR-5) [53, 54, 55]. Therefore, these receptors were selected for docking with the chimeric vaccine construct. The potential of chimeric vaccine construct docking with a Toll-Like Receptor was evaluated by the ClusPro 2.0 server (https://cluspro.bu.edu/login.php). ClusPro 2.0 is a rigid-body protein-protein docking service that predicts interactions between two proteins. This software is entirely automated and employs 3 distinct procedures: the first is a unique fast Fourier transform (FFT) correlation, the second is clustering the best energy conformations, and the third is evaluating cluster stability using brief Monte Carlo simulations [56, 57]. The PDB file of the refined vaccine construct was docked against TLR-1 (PDB Id: 6NIH), TLR-2 (PDB ID: 6NIG), TLR-4 (PDB ID: 3FXI), and TLR-5 (PDB ID: 3JOA) from the PDB database (https://www.rcsb.org/). The best-docked model was selected based on its lowest energy value among the models generated by the ClusPro 2.0 server.

### 2.16 Molecular dynamic simulation and protein-protein binding affinity analysis

To investigate the stability and dynamics of the docked complexes between MEV-TLR-1, MEV-TLR-2, MEV-TLR-4, and MEV-TLR-5, molecular dynamics simulations were conducted using the iMOD server (available at http://imods.chaconlab.org). This tool employs normal mode analysis (NMA) in internal coordinates to mimic the natural motions of biological macromolecules and generate plausible transition pathways between two similar structures, even with large molecules [58]. Additionally, the Prodigy web server, accessible at https://wenmr.science.uu.nl/prodigy/, was utilized to predict the binding affinity and dissociation constant of the docked proteins. Default parameters were applied for all analyses [59, 60, 61].

### 2.17 Immune simulation

The vaccine amino acid sequence was inputted into the C-ImmSim server, accessible at https://kraken.iac.rm.cnr.it/C-IMMSIM/, to analyze the immune response profile in a computational model. The C-ImmSim server assesses both humoral and cellular responses to the vaccine model [62] using a Position-specific scoring matrix (PSSM) and machine learning techniques. PSSM simulates various anatomical regions in mammals, including the Bone marrow (which stimulates hematopoietic stem cells and myeloid cell production), the thymus (where naive T cells are selected to prevent autoimmunity), and tertiary lymphatic organs, to mimic real-life immune responses. Immune stimulation was conducted by injecting the designed peptide vaccine, with three shots administered at four-week intervals (days 0, 28, and 56), translating to time steps at 1, 84, and 168 for each injection. Each time step represents an 8-hour interval, with the initial infusion administered at time zero. This prime-booster-booster strategy at 4-week intervals, was aimed to achieve a durable protective immune response. It was administrated with 336 simulation steps, with all other simulation parameters set to default values [19, 37].

### 2.18 Codon adaptation

Ensuring the efficacy of the vaccine model during cloning and expression is crucial for its in-vitro production. To achieve this, codon adaptation and *in-silico* cloning were carried out to express the final vaccine construct in a mammalian host. This step was necessary due to differences in codon usage between mammals and *Salmonella enterica* strains, aiming to ensure successful expression in the chosen host organism. The Java Codon Adaptation Tool (JCAT) server (http://www.prodoric.de/JCat) was utilized to perform reverse translation of the vaccine construct’s protein sequence into a DNA sequence. Attention was paid to avoid rho-independent transcription termination, prokaryotic ribosome binding sites, and restriction enzyme cleavage sites [63]. In JCAT, a codon adaptation index (CAI) score of 1.0 is considered ideal, although scores above 0.8 are generally satisfactory. Additionally, the sequence’s GC content was optimized to fall within the favorable range of 30% to 70%.

### 2.19 Restriction enzyme mapping and *in-silico* cloning into plasmid

To analyze restriction enzyme sites within the JCAT codon-optimized sequence of the vaccine construct, we utilized the NEBcutter Version 3.0.17 software (available at https://nc3.neb.com/NEBcutter/). NEBcutter V3.0 is specifically designed to support users in planning restriction digests and molecular cloning projects. By submitting a sequence, or selecting from a library of common plasmids, users can generate visual maps of restriction enzyme sites and simulate digests with chosen enzymes. In our study, we introduced HindIII and BamHI restriction sites at the N-terminal and C-terminal regions of the DNA sequence respectively, as optimized by the JCAT server. Additionally, a hexa histidine tag (6xHis tag) was incorporated just before the BamHI site at the C-terminal to facilitate protein purification. For the cloning process, we employed the SnapGene software to insert our DNA sequence into the pcDNA3.1_CT-GFP expression vector (detailed at https://www.snapgene.com/plasmids/mammalian_expression_vectors/pcDNA3.1_CT-GFP). This vector, optimized for mammalian expression, was selected for its efficacy in visualizing transfection in mammalian cell lines [48, 63, 64].

## 3.0 Results

### 3.1 Retrieval of *S.* Kentucky OMPs sequences and generation of consensus sequence

The sequences for *Salmonella* Kentucky outer membrane proteins (Omps) A, C, and F, along with their respective accession numbers, were sourced from the NCBI protein database in FASTA format. A total of 65 outer membrane proteins (Omps) of *Salmonella* Kentucky were retrieved, comprising of 18 OmpA, 32 OmpC, and 15 OmpF (Table 1). The FASTA format sequences of all 65 OMPs A, C, and F classes were aligned separately by Jalview 2.11.2.7. Additionally, the FASTA sequence of *fliC* protein (GenBank accession number KAA0422324.1) of *Salmonella* Typhimurium used as an Adjuvant was also retrieved (Table 2).

**Table 1.**
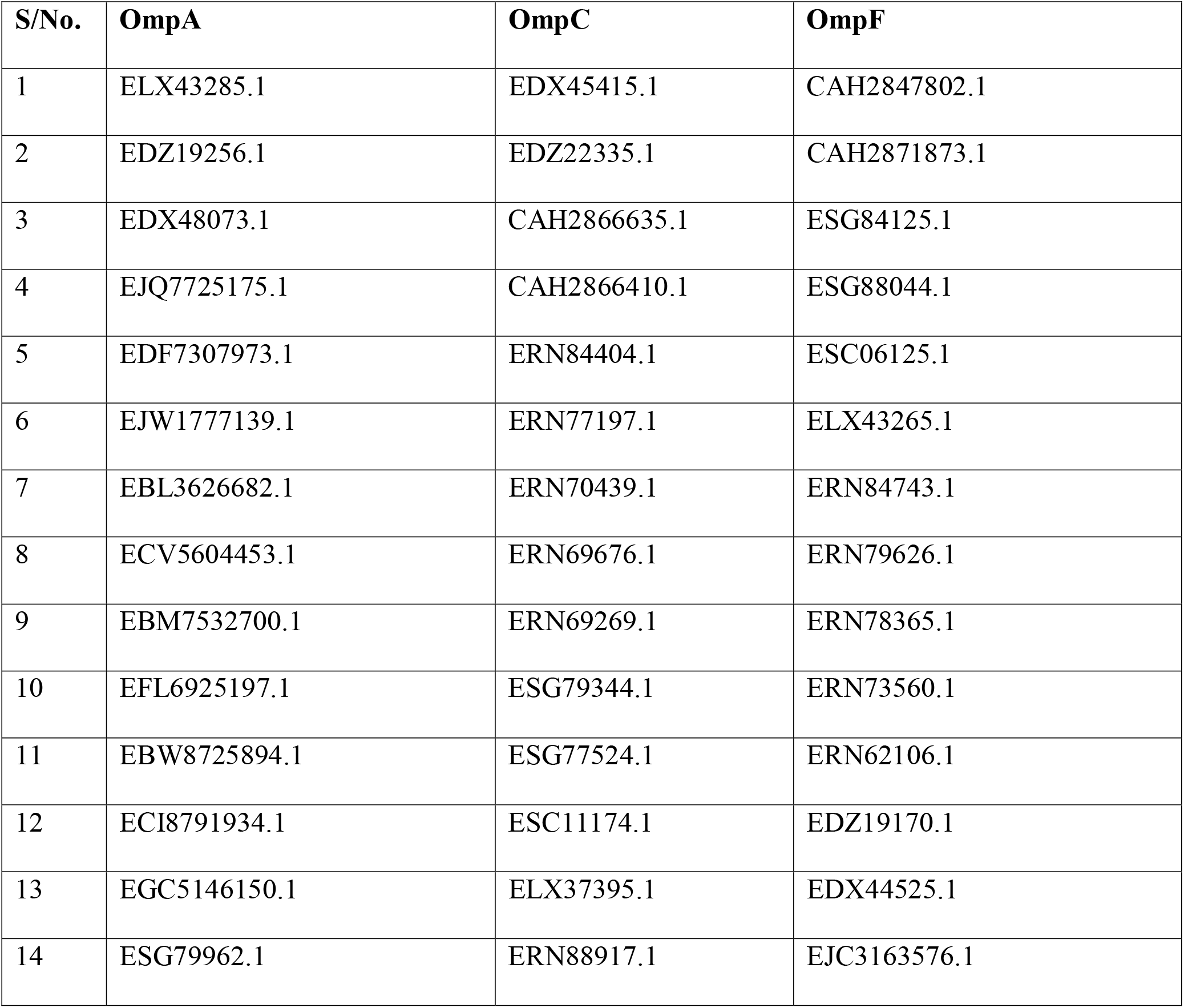
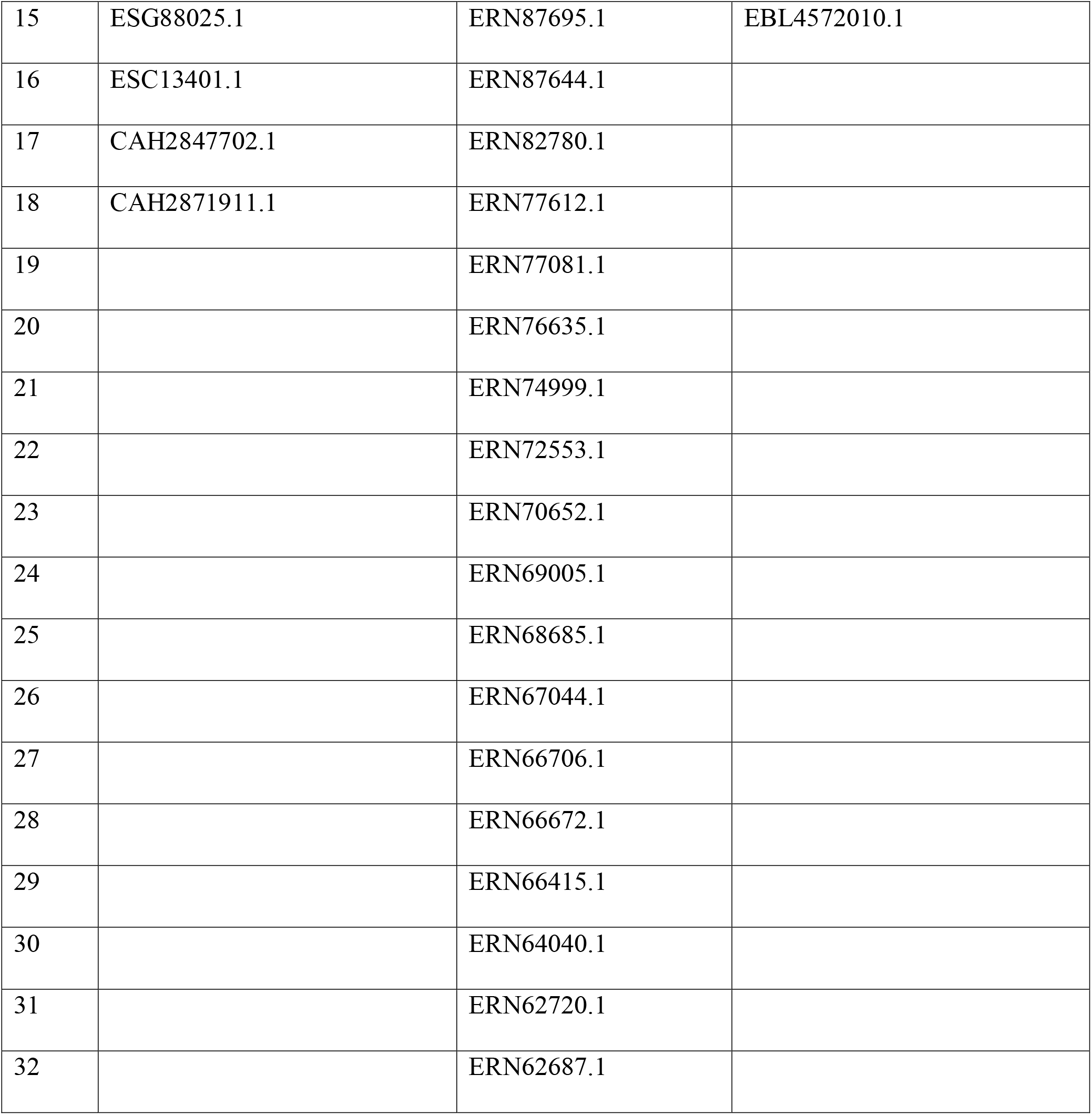
The accession number of the outer membrane proteins A, C, and F selected for the development of MEV was retrieved from the NCBI protein database.

**Table 2.**
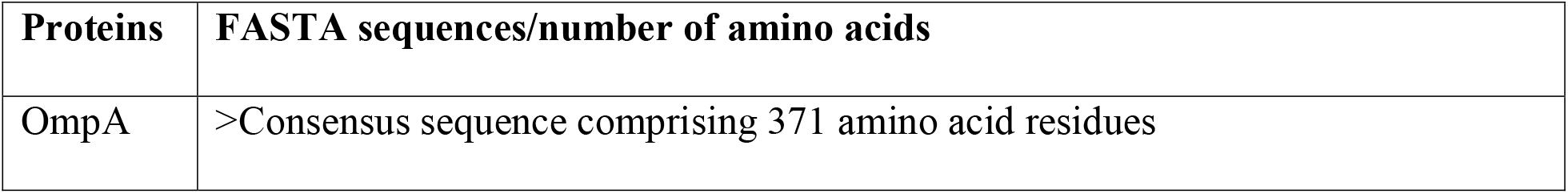
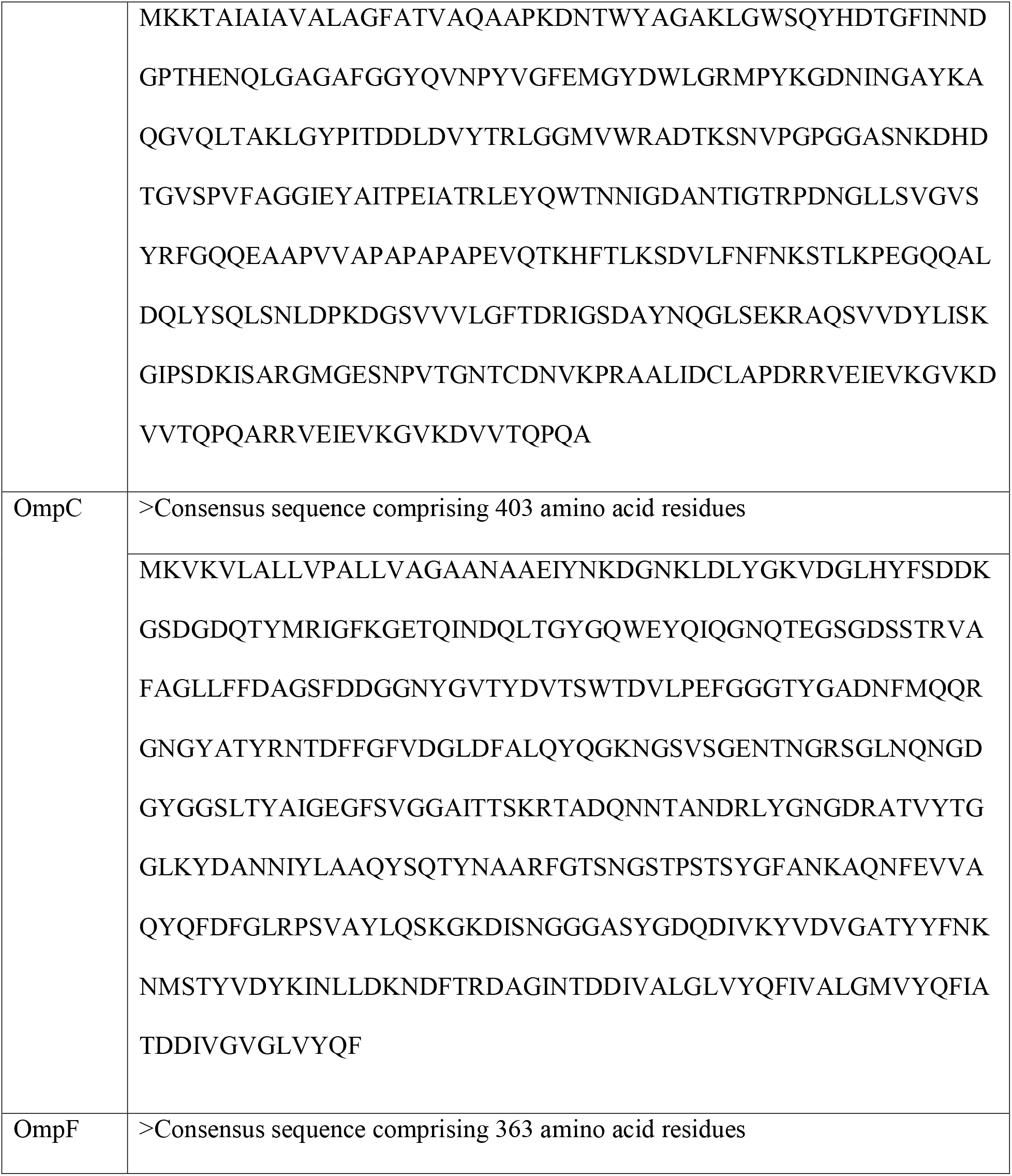
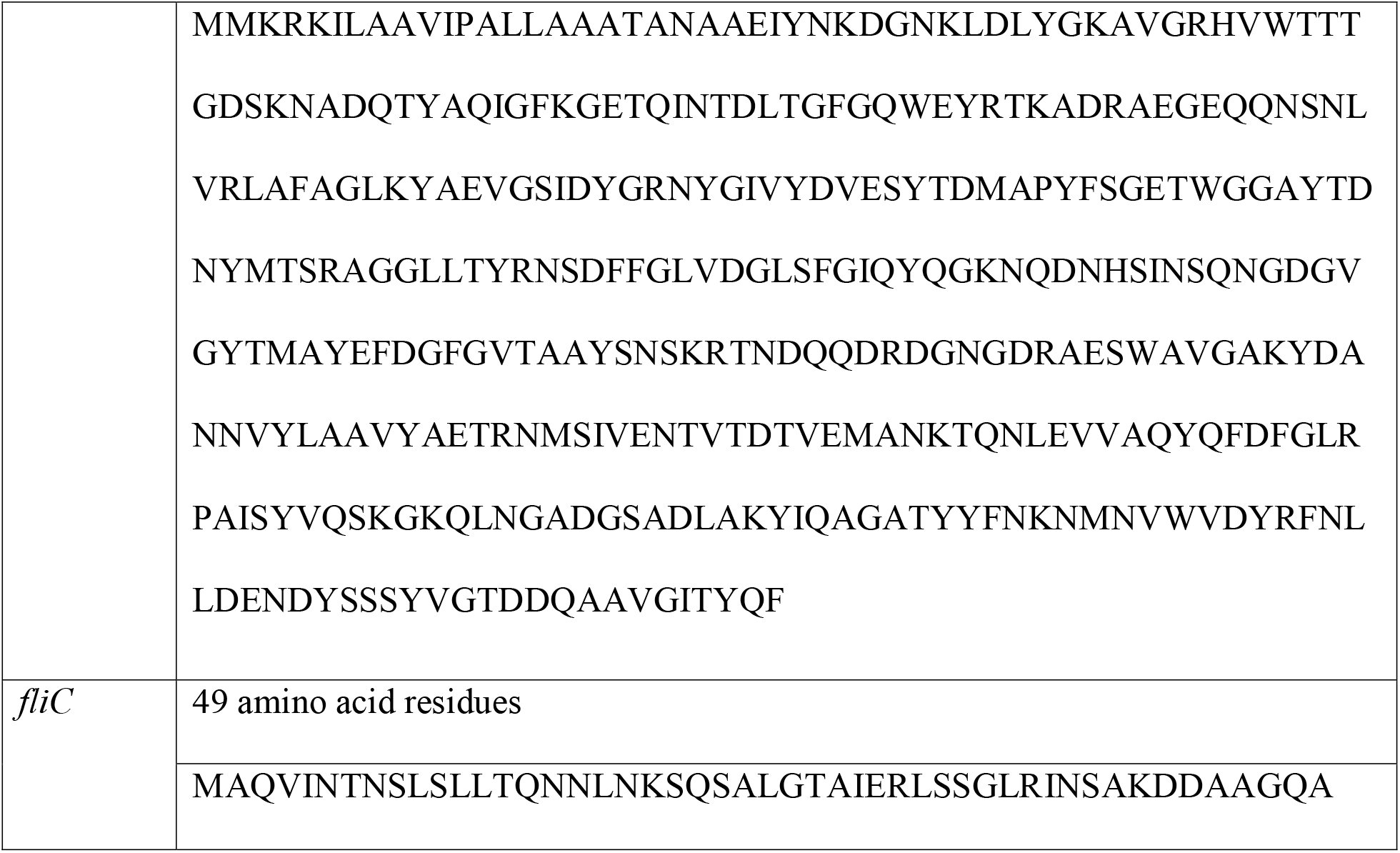
Jalview version 2.11.2.7 generated consensus sequences of the outer membrane proteins A, C, and F and selected *fliC* protein for the development of MEV.

### 3.2 Results for CTL and HTL epitopes prediction and epitope conservancy analysis

At a threshold value of 0.5 percent for strong binders and following antigenicity, allergenicity, and toxicity screening, a final total of 4 CTL epitopes for OmpA, 5 CTL epitopes for OmpC, and 5 CTL epitopes for OmpF were selected for the multi-epitope vaccine construction (Table 3). Their corresponding allele(s) and positions were noted (Table 4). The result from the IEDB server using the seven-allele reference set and applicable thresholds generated 4, 2, and 2 HTL epitopes for OmpA, OmpC, and OmpF respectively. These made up the HTL epitopes selected for constructing the MEV (Table 3). Their corresponding allele(s) and positions were noted (Table 4). Also, the IEDB server conservancy analysis results at > = 100 % protein sequence identity threshold predicted a minimum and maximum identity of 100 % for all epitopes across the different classes of OMPs A, C, and F CTLs and HTLs. Epitope “IEYAITPEI” had a 100 % minimum and maximum identity conservancy as an OmpA, but had a minimum identity at 33.33 % and a maximum identity at 33.33 % as an OmpC CTL epitope (Table 4).

**Table 3.**
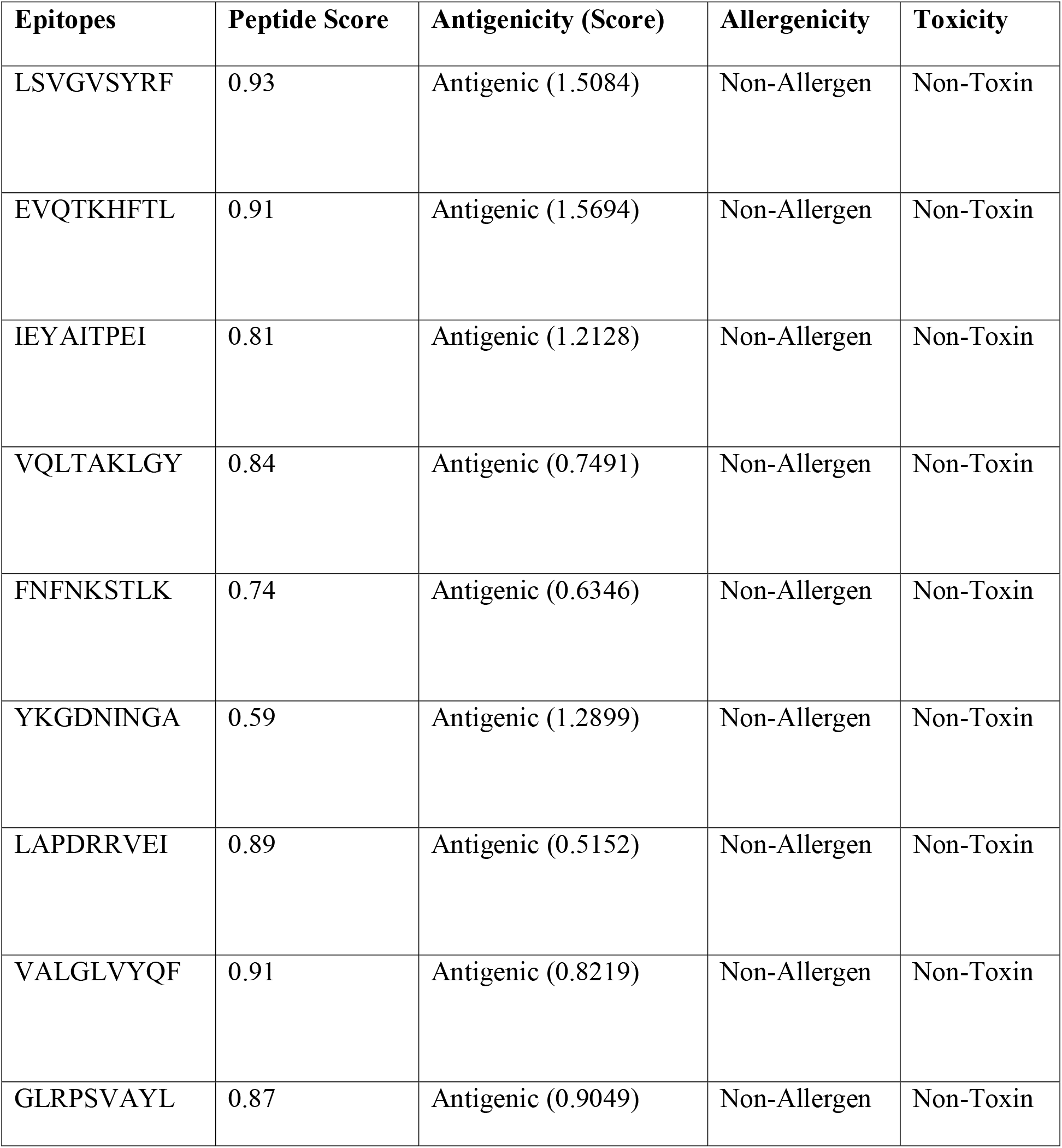
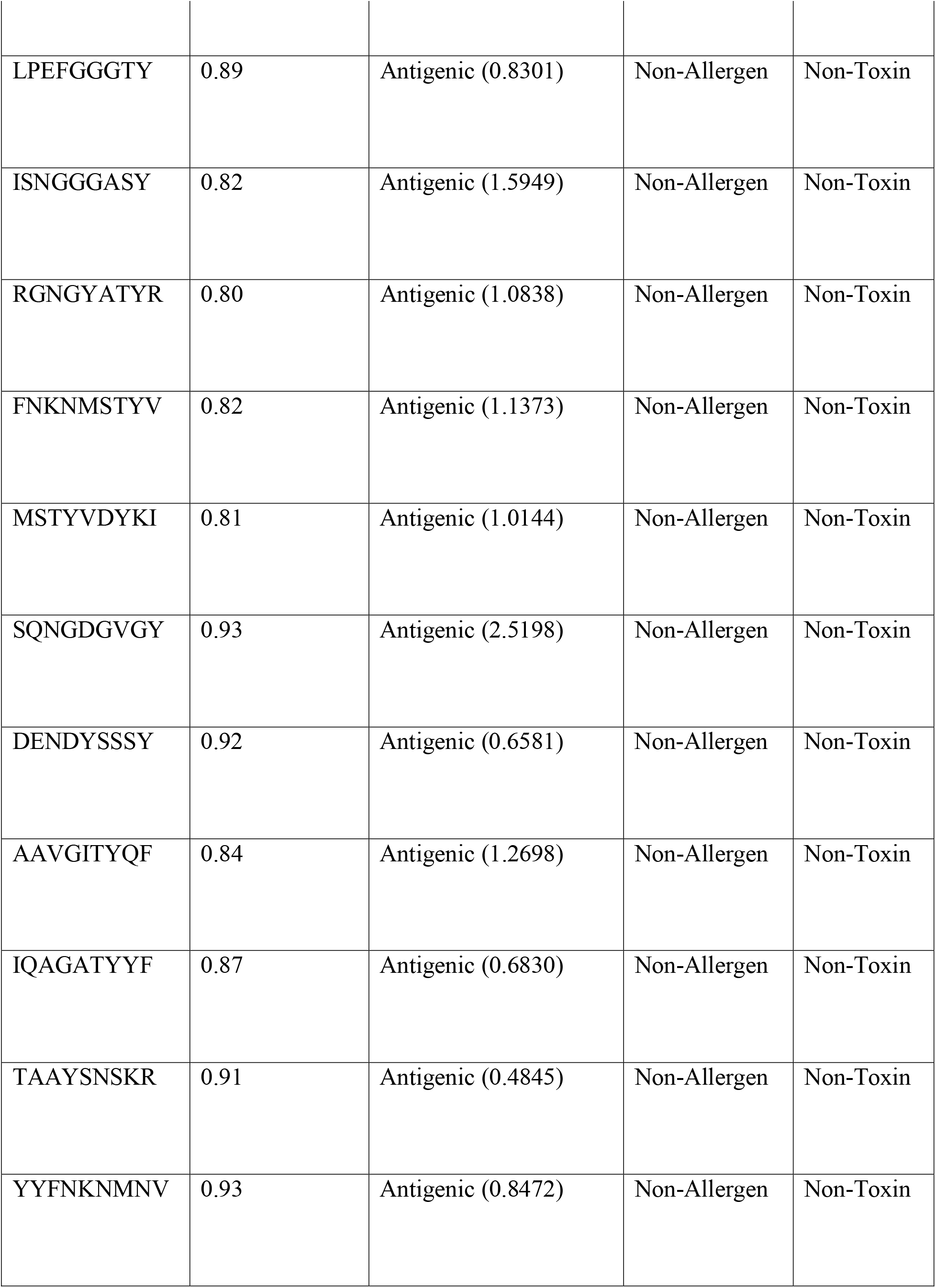
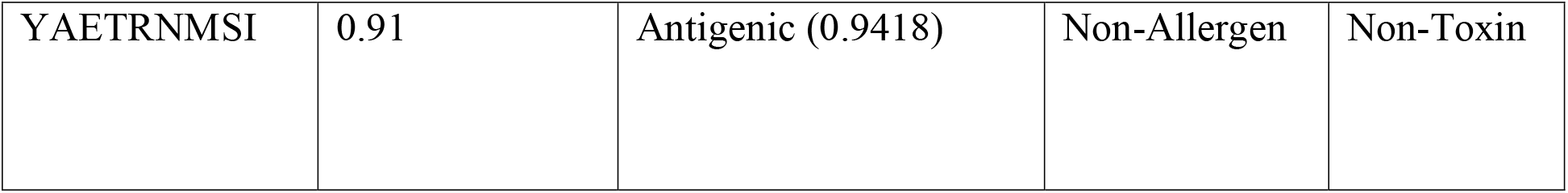
Predicted CTL and HTL epitopes of *S.* Kentucky OmpA, OmpC, and OmpF proteins from consensus sequences with their immunogenic properties.

**Table 4.**
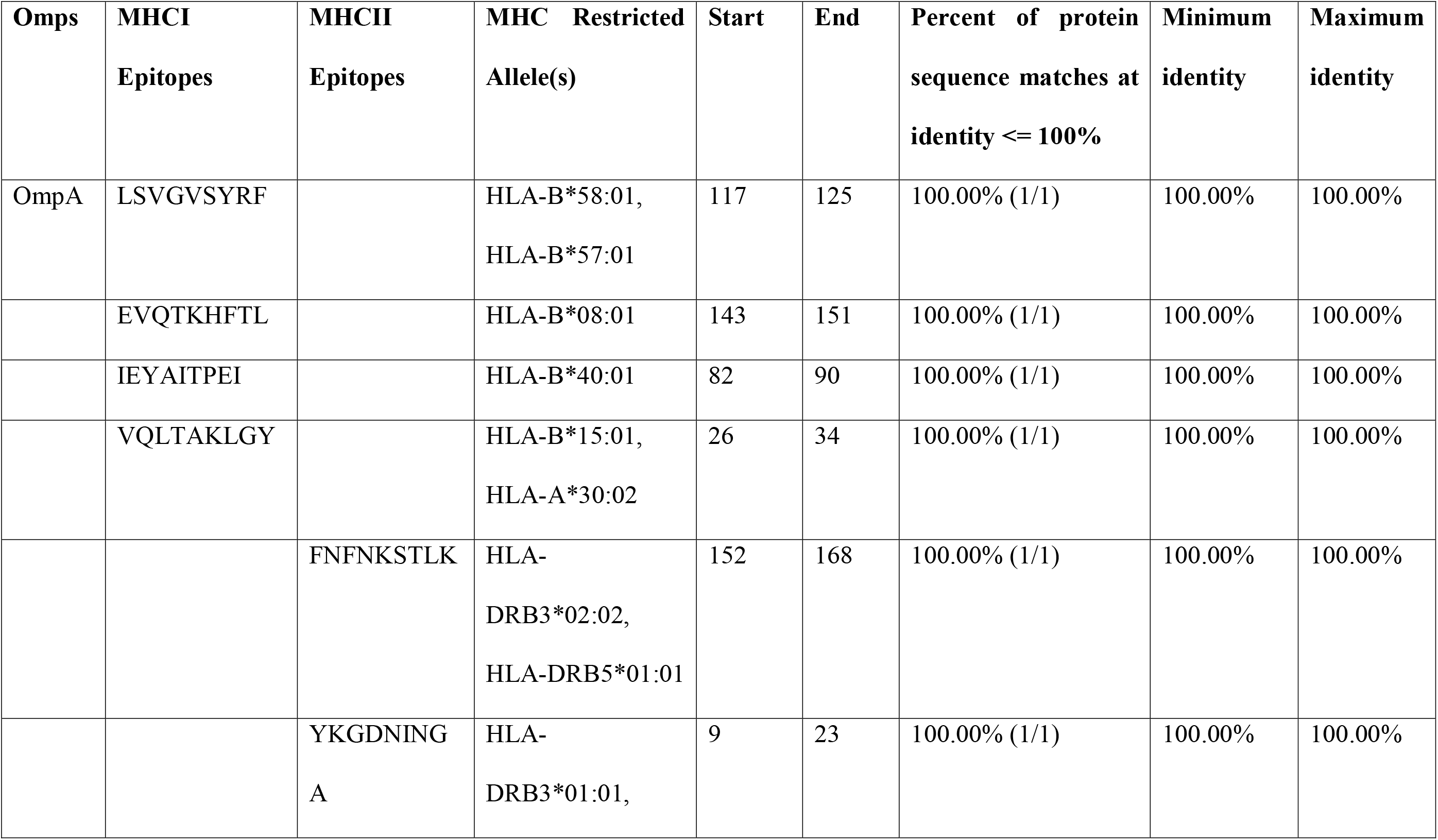
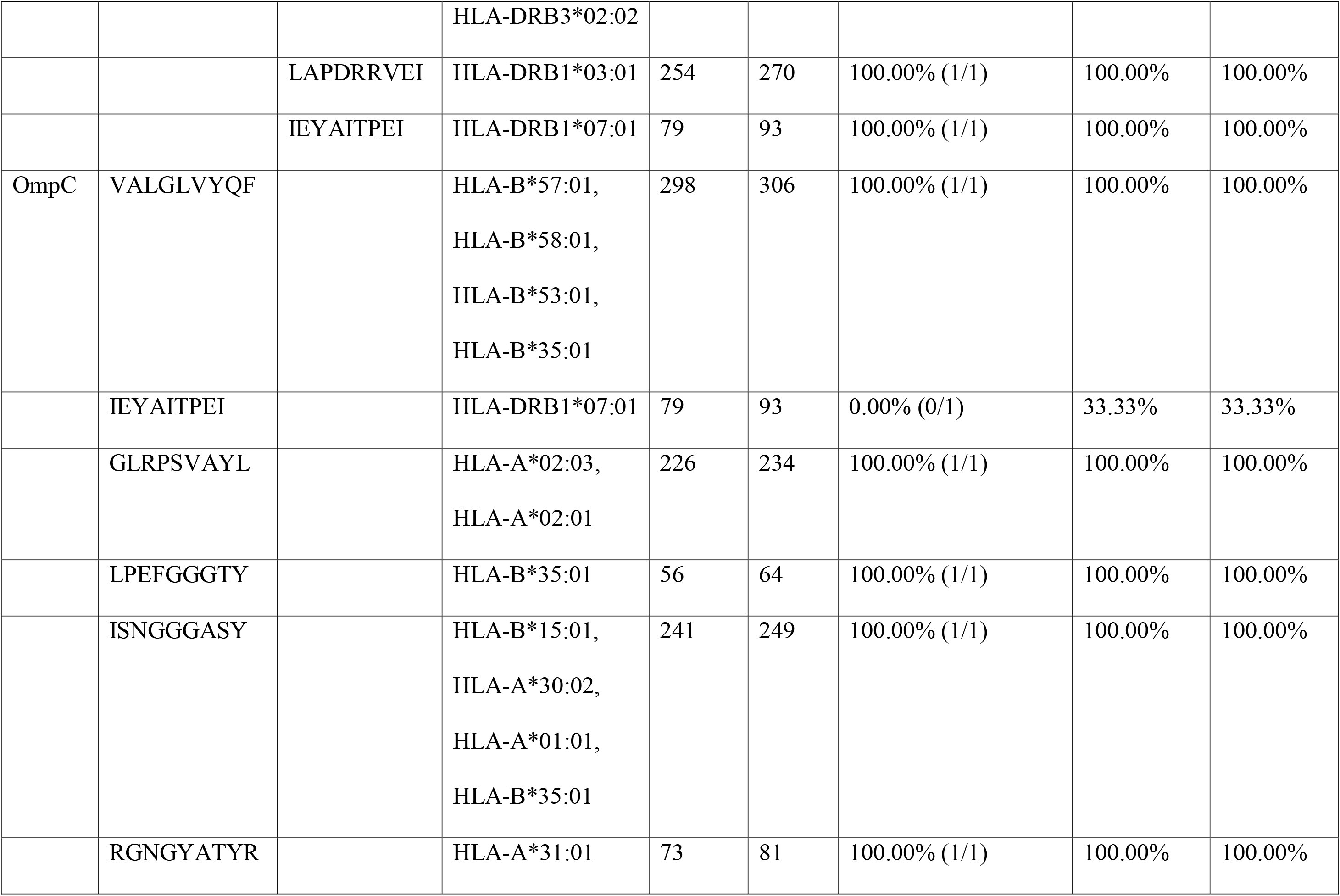
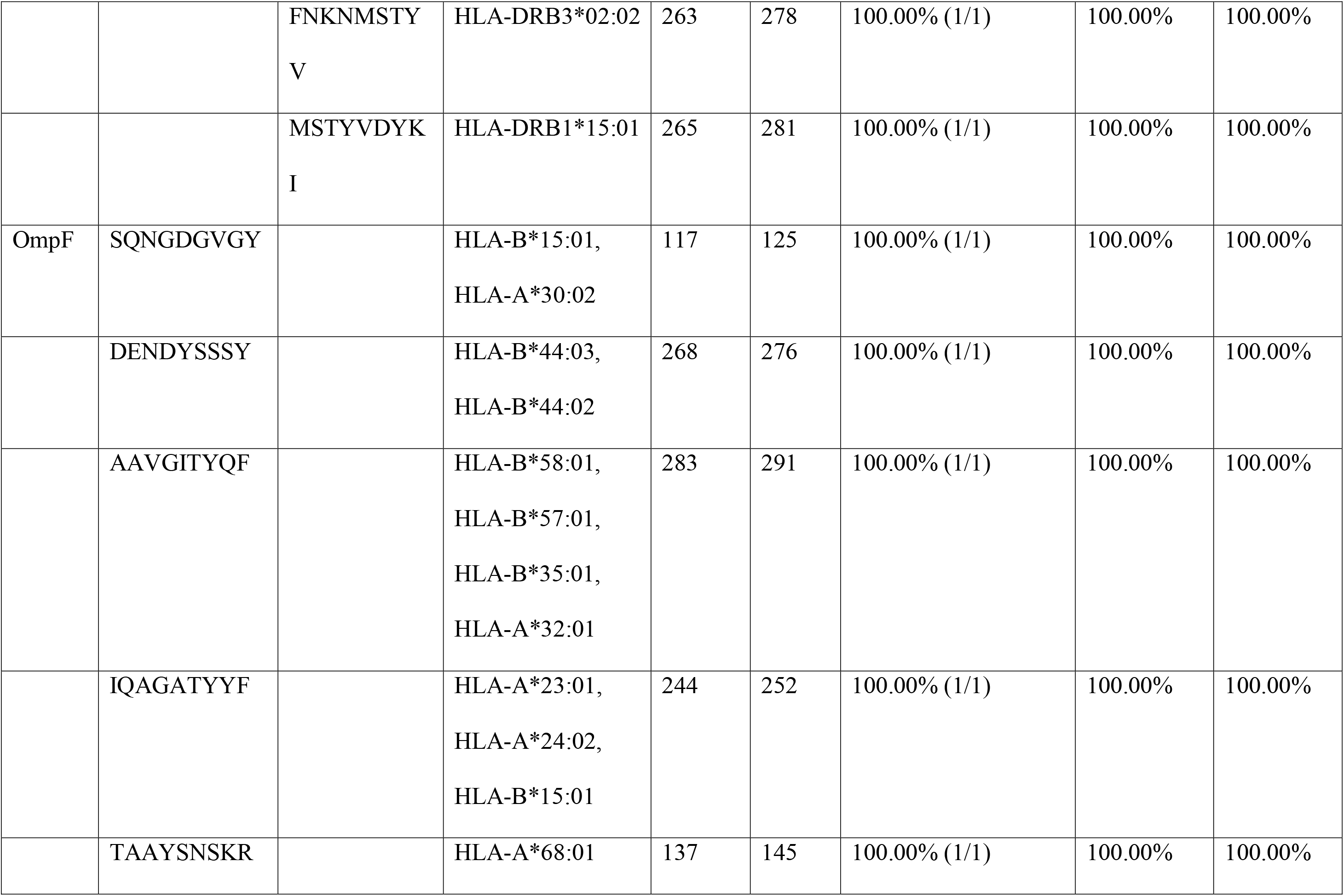
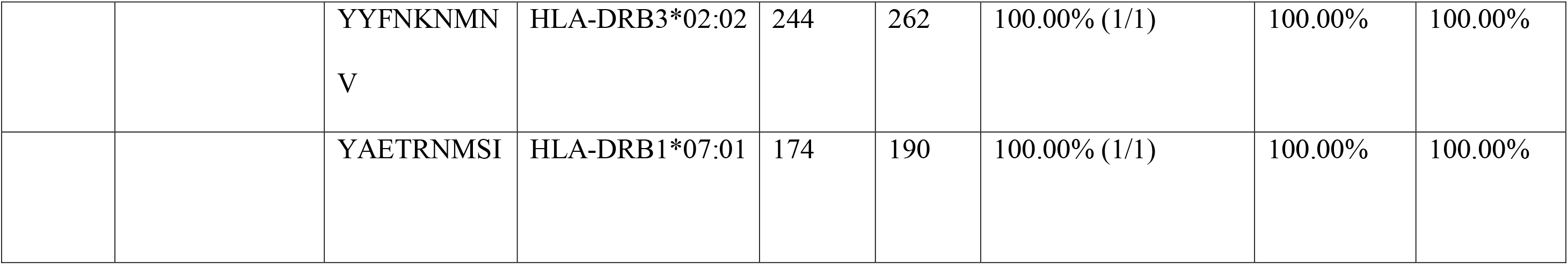
The predicted MHC-restricted allele(s) of the CTL and HTL epitopes and degree of conservancy for the OMPs A, C, and F screened for antigenicity, allergenicity, and toxicity.

### 3.3 Linear B-lymphocytes epitopes prediction

ABCpred server predicted a total of 35 B-cell epitopes for the consensus sequence of OmpA, 39 B-cell epitopes for the consensus sequence of OmpC and 41 B-cell epitopes for the consensus sequence of OmpF with a cut off binding score at 0.51. B-cell epitopes with a high score of ≥ 0.85 and a minimum peptide rank score of 7 were further subjected to antigenicity, allergenicity and toxicity screening. Finally, 2 B-cell epitopes for OmpA, 2 B-cell epitopes for OmpC and 3 B-cell epitopes for OmpF were selected for the MEV construction (Table 5).

**Table 5.**
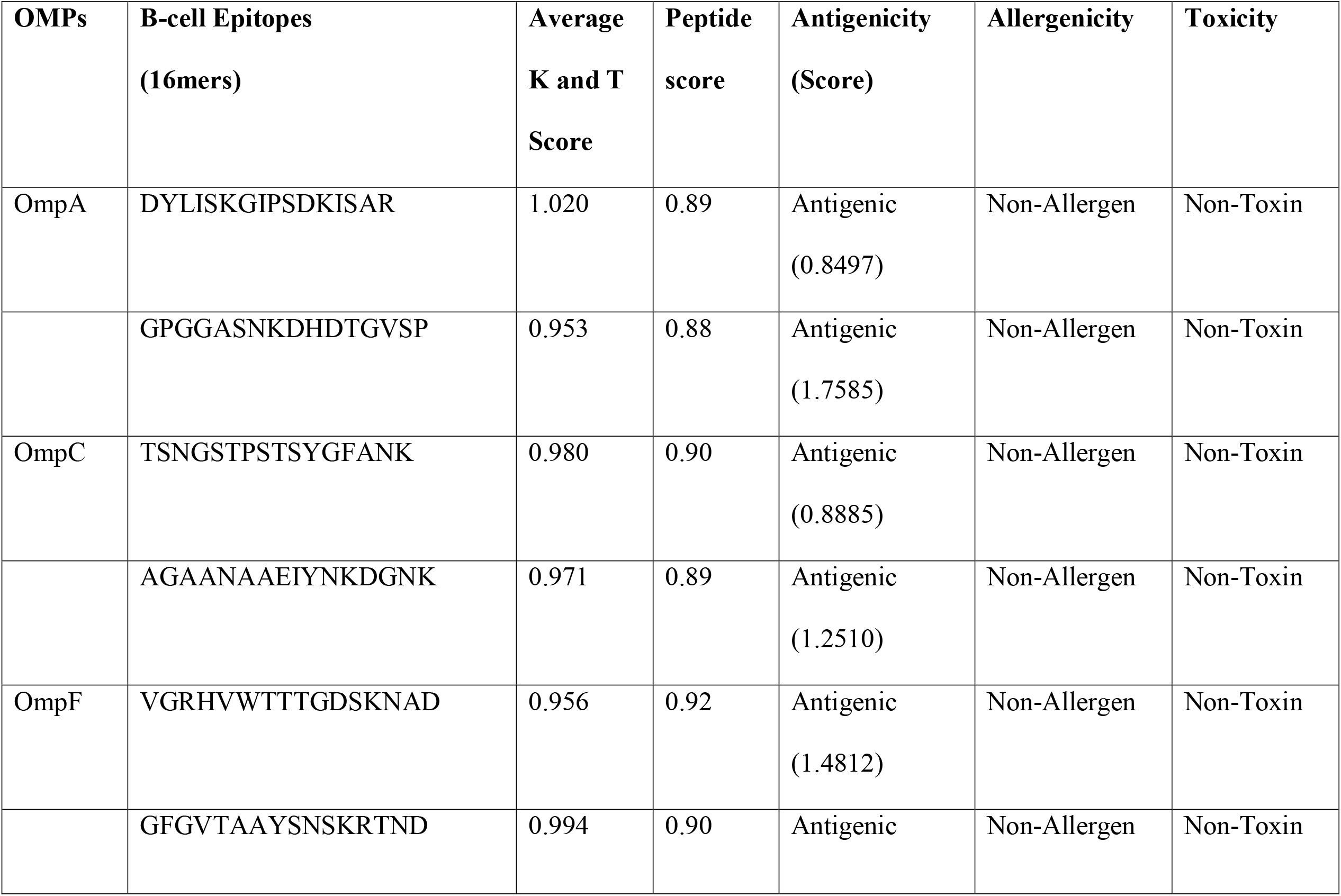
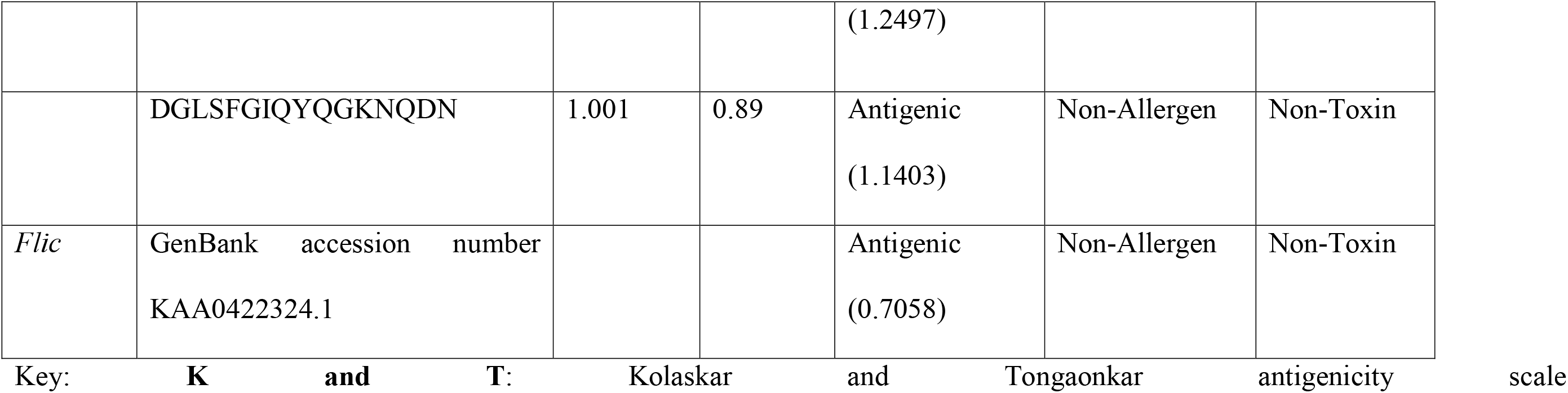
Predicted linear B-cells epitopes of *S*. Kentucky OMP A, C and F from consensus sequences and *flic* protein (adjuvant) with their immunogenic properties.

### 3.4 Interferon gamma (IFN-γ) inducing epitope prediction

Eight MHC class II epitopes were submitted to the IFNepitope server for IFN-γ prediction. Two HTL epitopes were positive, thus; capable of inducing the production of interferon gamma. The SVM method predicted epitope “LAPDRRVEI” as a positive inducer and the MERCI method predicted “IEYAITPEI” epitope as a positive inducer (Table 6).

**Table 6.**
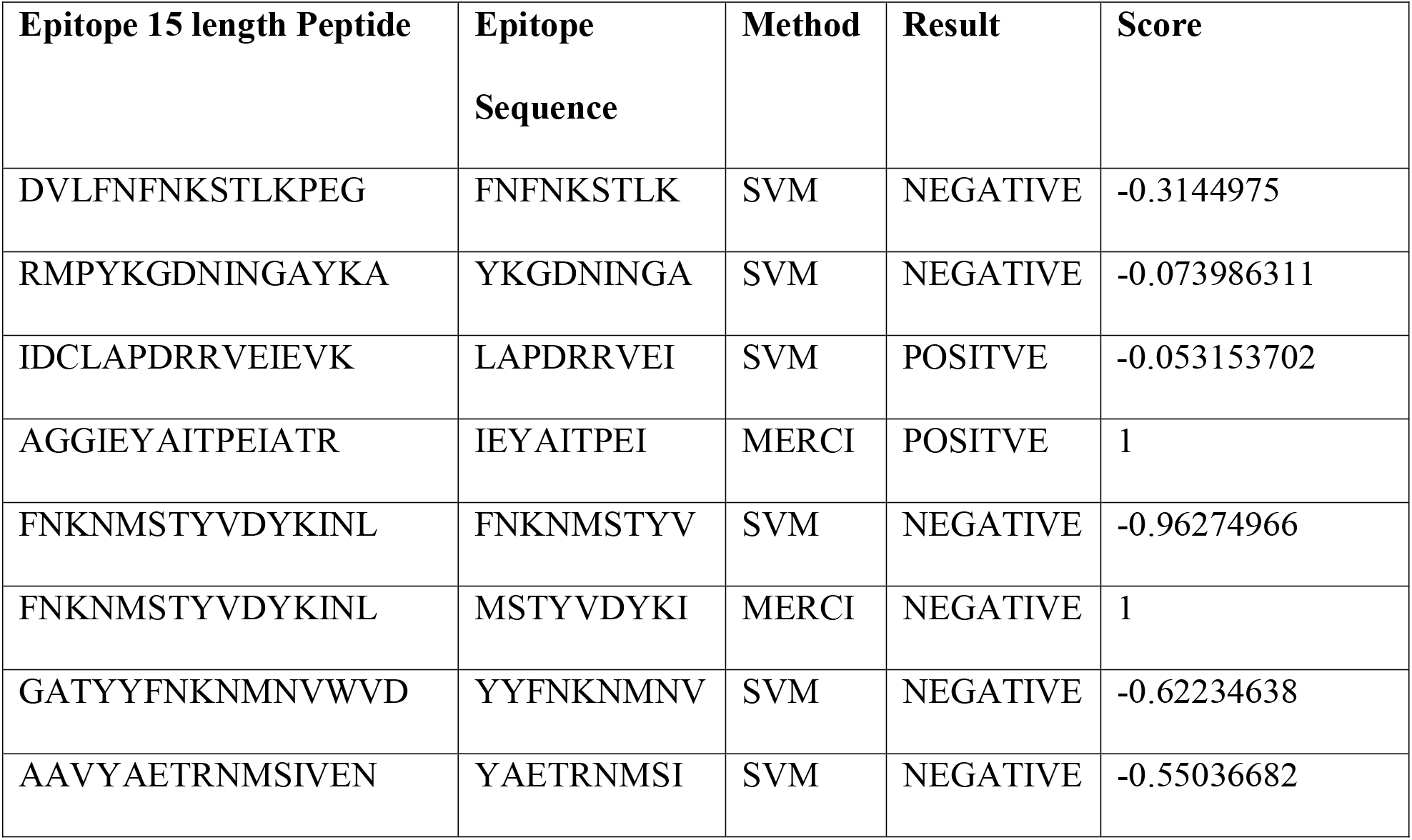
Predicted OMPs A, C and F MHC Class II T-cell epitopes capable of inducing interferon gamma (IFN-γ)

### 3.5 Population coverage

The worldwide population percentage coverage of the selected putative HLA class I and HLA class II restricted epitopes was 94.91 %. This analysis of the MHC-I and MHC-II alleles confirmed that the world population will be covered if the intended vaccine architecture was administered. Based on continental and regional analysis, Europe, North America and North Africa had the highest population coverage of 97.59 %, 96.92 % and 92.89 % respectively (Fig 1). On the African continent, the percentage population coverage for West Africa, South Africa, East Africa and Central Africa were 89.77 %, 83.02 %, 81.86 % and 78.17 % respectively with a class combined regional average of 85.14 (Table 7). Outside the African continent, East Asia, North-East Asia and South America were 90.82 %, 79.71 % and 83.0 % respectively (Fig 1). Furthermore the selected putative HLA class I restricted epitopes exhibited a higher individual percentage coverage (90.02 %) when queried against the HLA class II restricted epitopes (49.02 %) across the world (Fig 1).

**Fig 1.**
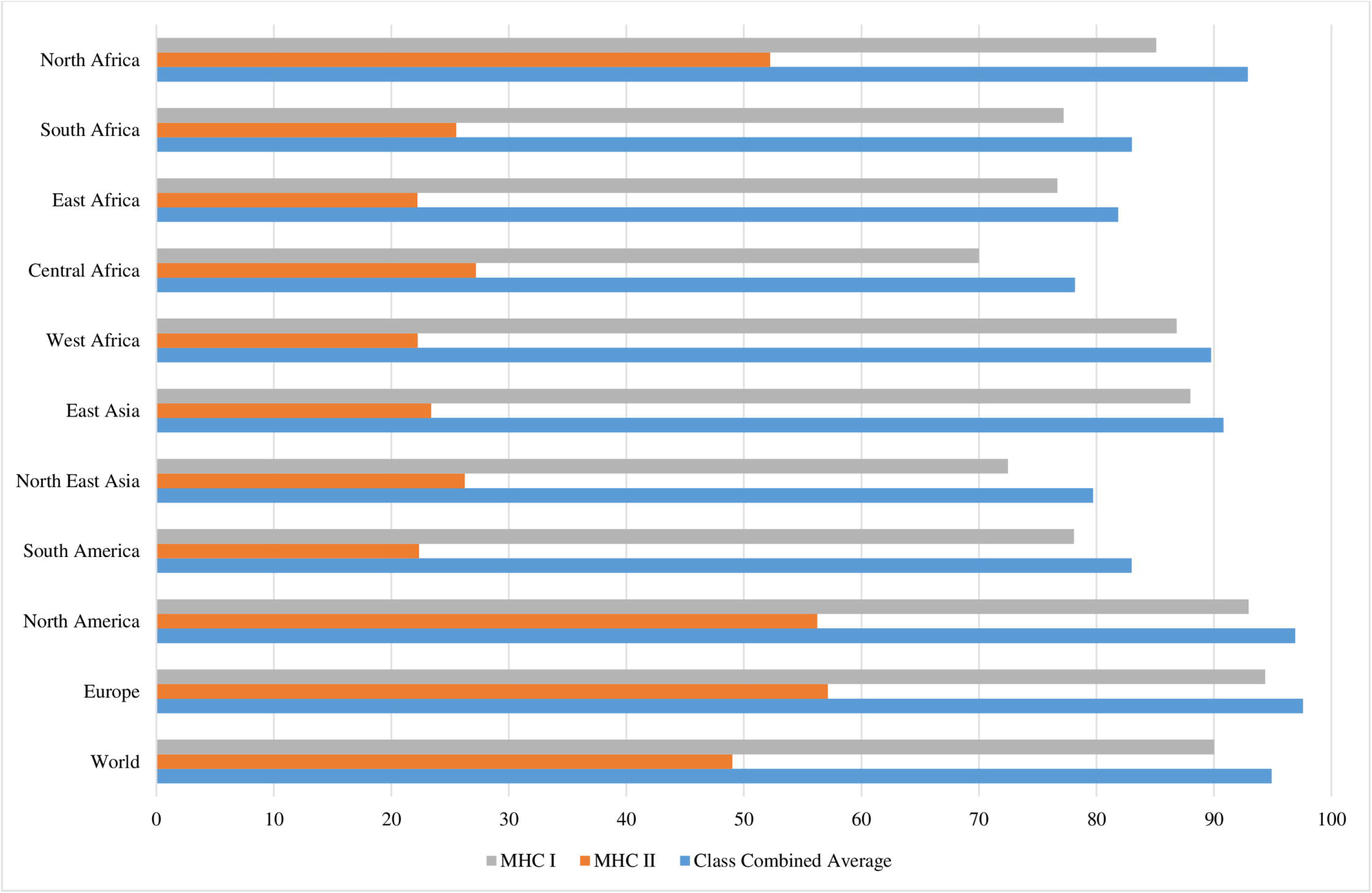
Continental and regional coverage of the MHC class I and MHC class II T-Cell epitopes.

### 3.6 Construction of MEV sequence and interferon gamma (IFN-γ) induction capabilities of the MEV

The MEV construct consists of 462 amino acids residues from a total of 28 selected antigenic T-cells and B-cell epitopes comprising 14 CTLS, 7 HTLs and 7 B-cell epitopes, covalently linked to an immuno-adjuvant (*Salmonella* Typhimurium *fliC* protein; GenBank accession number KAA0422324.1 with 49 amino acid residues). The *fliC* protein was fused to the epitopes of CTLs with an EAAAK linker at the N-terminal end of the vaccine construct. The CTL peptides of OmpA, OmpC and OmpF were fused together with an AAY linker and fused to the first HTL OmpA epitope with an AAY linker. The HTL peptides and B-cell epitopes were fused together with a GPGPG linker with the B-cell epitope of OmpF at the C-terminal end (Fig 2). Result of antigenicity, allergenicity and toxicity prediction on the MEV construct confirms the vaccine as antigenic (with a score of 1.0458), non-allergenic and non-toxin.

**Fig 2.**
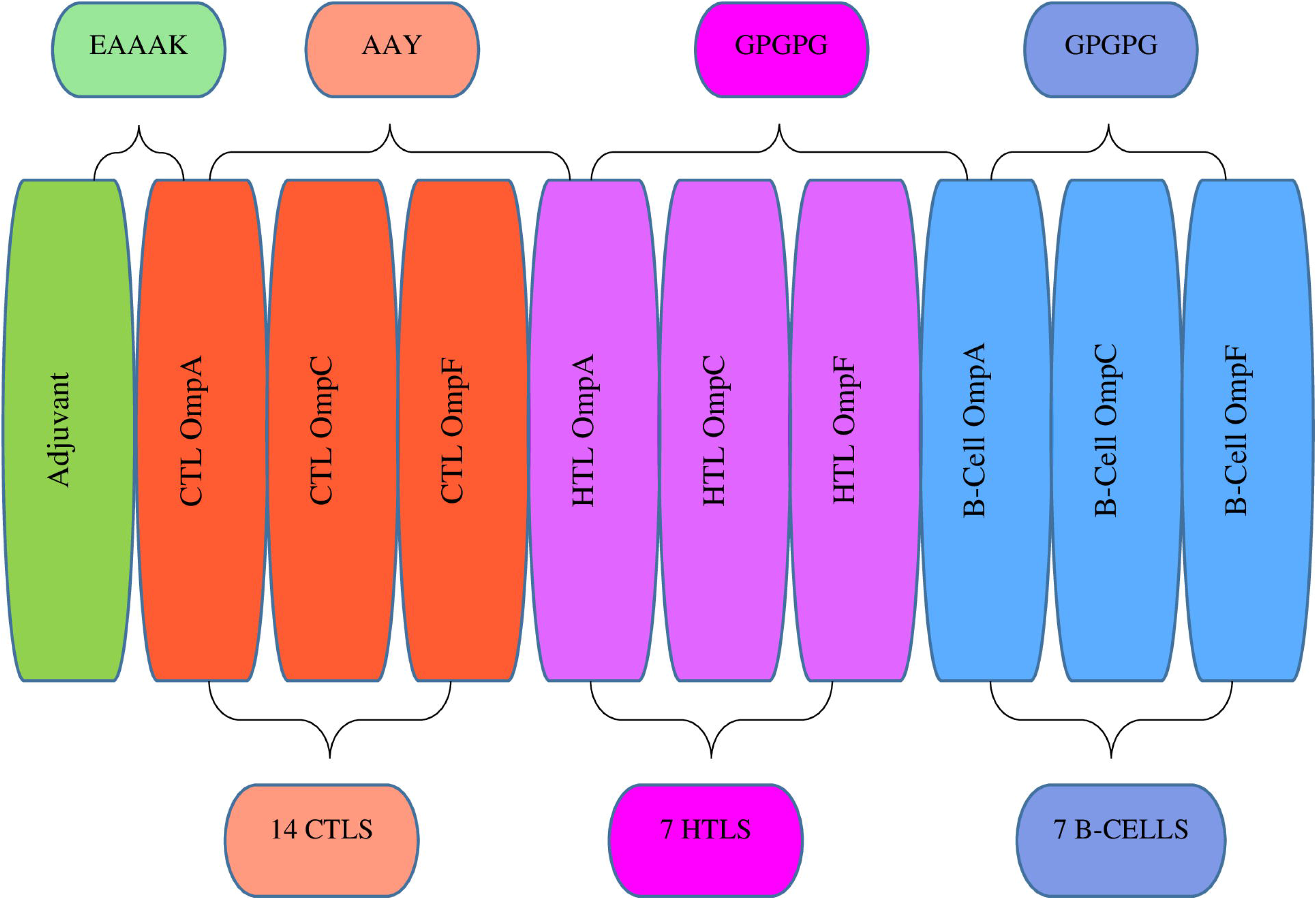
The schematic structural arrangement of the final multiepitope vaccine construct.

The IFNepitope server support vector machine (SVM) hybrid model result confirmed a positive result capable of inducing IFN-γ and a score of 1.5695871.

### 3.7 Prediction of physicochemical characteristics and solubility of the MEV

The physicochemical parameters and solubility properties of the chimeric vaccine construct using Expasy Protparam and the Protein-Sol servers expressed the following. The molecular weight of the vaccine construct was 47343.98 Da and the bio-computed theoretical isoelectric point (pI) was 9.150. The estimated half-life in mammalian cell is 30 hours, >20 hours and >10 hours in yeast and *E. coli* respectively. The solubility of the vaccine was found to be 0.452 on the protein sol server (value > 0.45 is predicted to have a higher solubility than the average soluble *E. coli* protein). The instability index of the vaccine was 25.66, signifying that the vaccine is stable in a solvent environment (*>* 40 signifies instability). The computed aliphatic index of the protein is 57.23. The aliphatic index quantifies the relative volume occupied by the aliphatic side chains of specific amino acids: alanine, valine, leucine, and isoleucine. A higher aliphatic index is associated with increased thermo-stability in globular proteins. The GRAVY score for the vaccine proteins was calculated to be −0.429, indicating they have good interactions with water molecules. The general average hydropathicity (GRAVY) score is a common parameter used in codon usage bias (CUB) analysis to assess a protein’s hydrophobic and hydrophilic properties. The GRAVY score ranges from −2 to 2, where positive values indicate hydrophobic proteins and negative values indicate hydrophilic proteins.

### 3.8 Prediction of secondary and tertiary structure

The result for secondary structure analysis of the designed MEV from PSIPRED and SOPMA server indicated that 77 amino acids (16.67 %) were alpha-helices, 41 amino acids (8.87 %) were extended beta strands, 21 amino acids (4.55 %) were beta turns, and 323 amino acids (69.91 %) were random coils (Fig 3). The 3D structure was built by the I-Tasser server. According to the threading templates, 5 tertiary 3D structures were predicted. The best C-score model with a −0.69 C-score value was selected for further refinement and validation. The consistency of the 3D model was improved by the Galaxy refine web server. Five improved refined models were produced from the initial MEV model (the −0.69 C-score structure from I-Tasser). Model 2 was the most significant model based on the computational structural qualities. Model 2 had a GDT-HA score of 0.9221, RMSD of 0.493, MolProbity of 2.352, a Clash score of 13.7, a zero Poor rotamers (0.0) and Rama favoured value of 82.0 (Fig 4).

**Fig 3.**
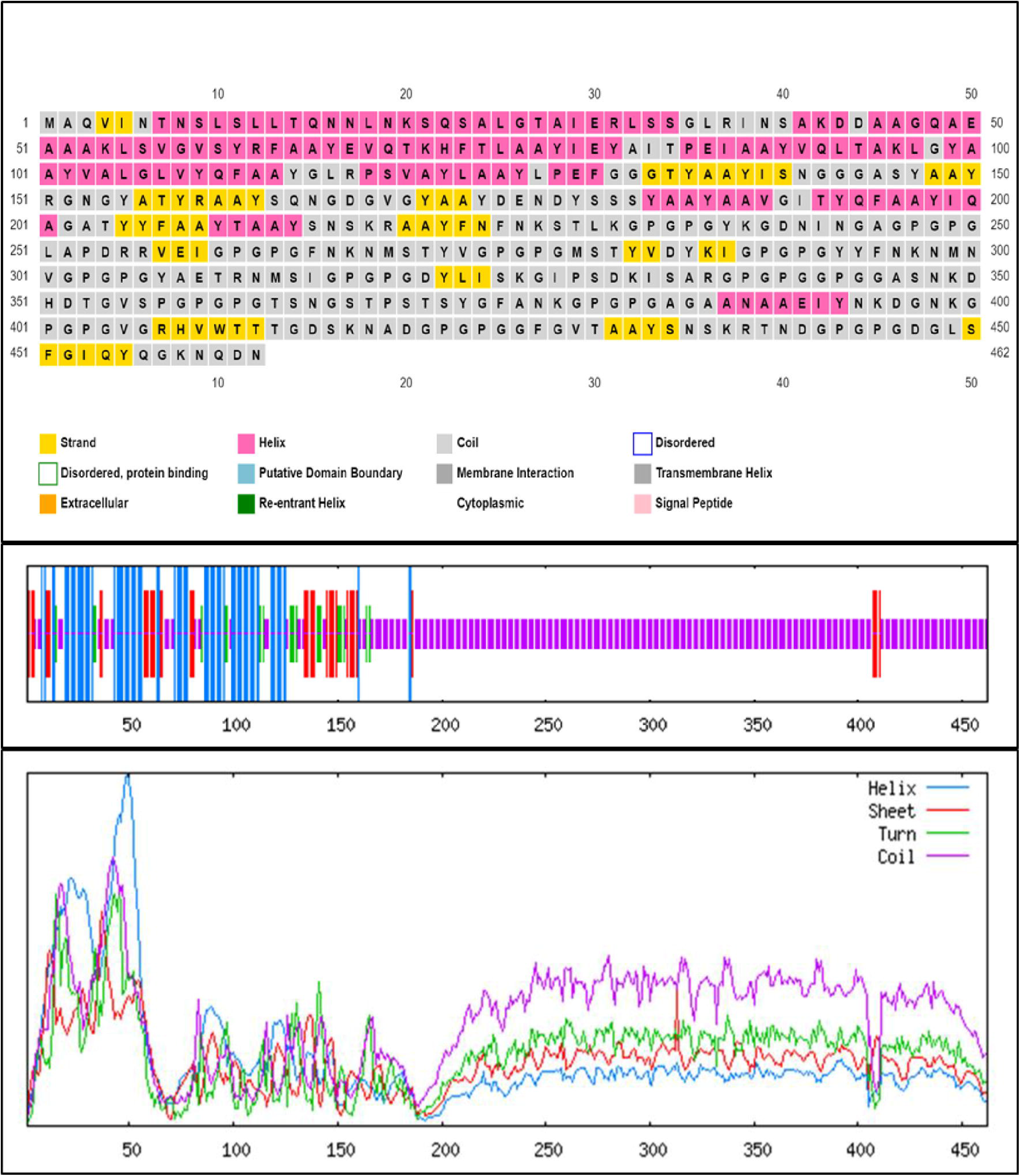
The PSIPRED and SOPMA server graphical demonstration of secondary structure properties of the final designed MEV.

**Fig 4.**
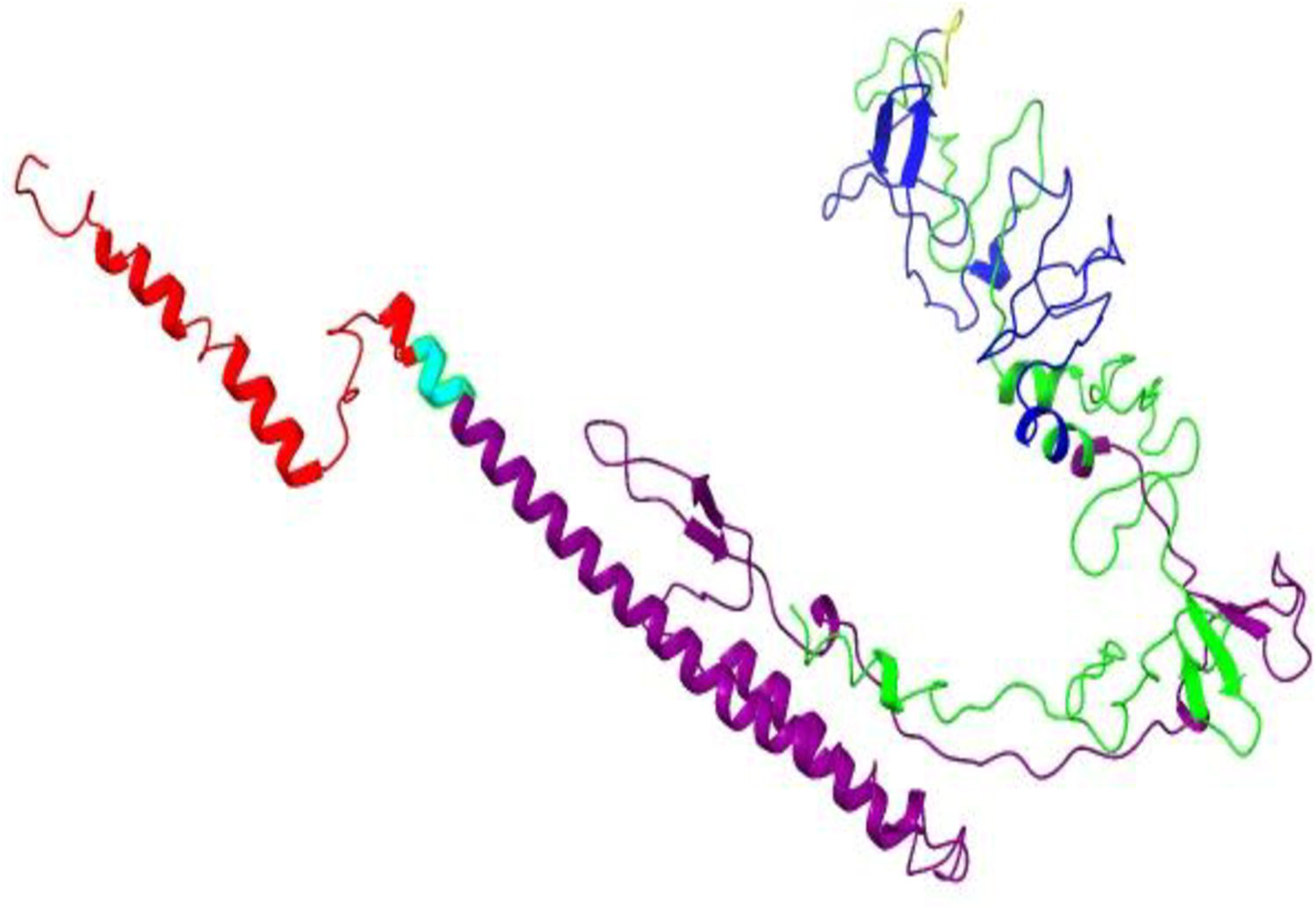
The tertiary structure of the designed MEV was predicted by I-TASSER and coloured with ChimeraX software. *fliC* protein (adjuvant) coloured in Red, CTLs in Purple, HTLs in Blue, and B-cell epitopes in Lime green

The validation of the refined tertiary structure of the MEV on the PROCHECK and VADAR version 1.8 servers produced a Ramachandran plot with 264 residues (75.6 %) in the most favored region, 57 residues (16.3%) in additional allowed regions, 19 residues (5.4 %) in generously allowed regions and 9 number of residues (2.6 %) in the disallowed regions, making a total of 349 number of non-glycine and non-proline residues. Additionally, there were 77 glycine residues, 34 proline residues, and 2 number of end-residues (excl. Gly and Pro) (Fig 5).

**Fig 5.**
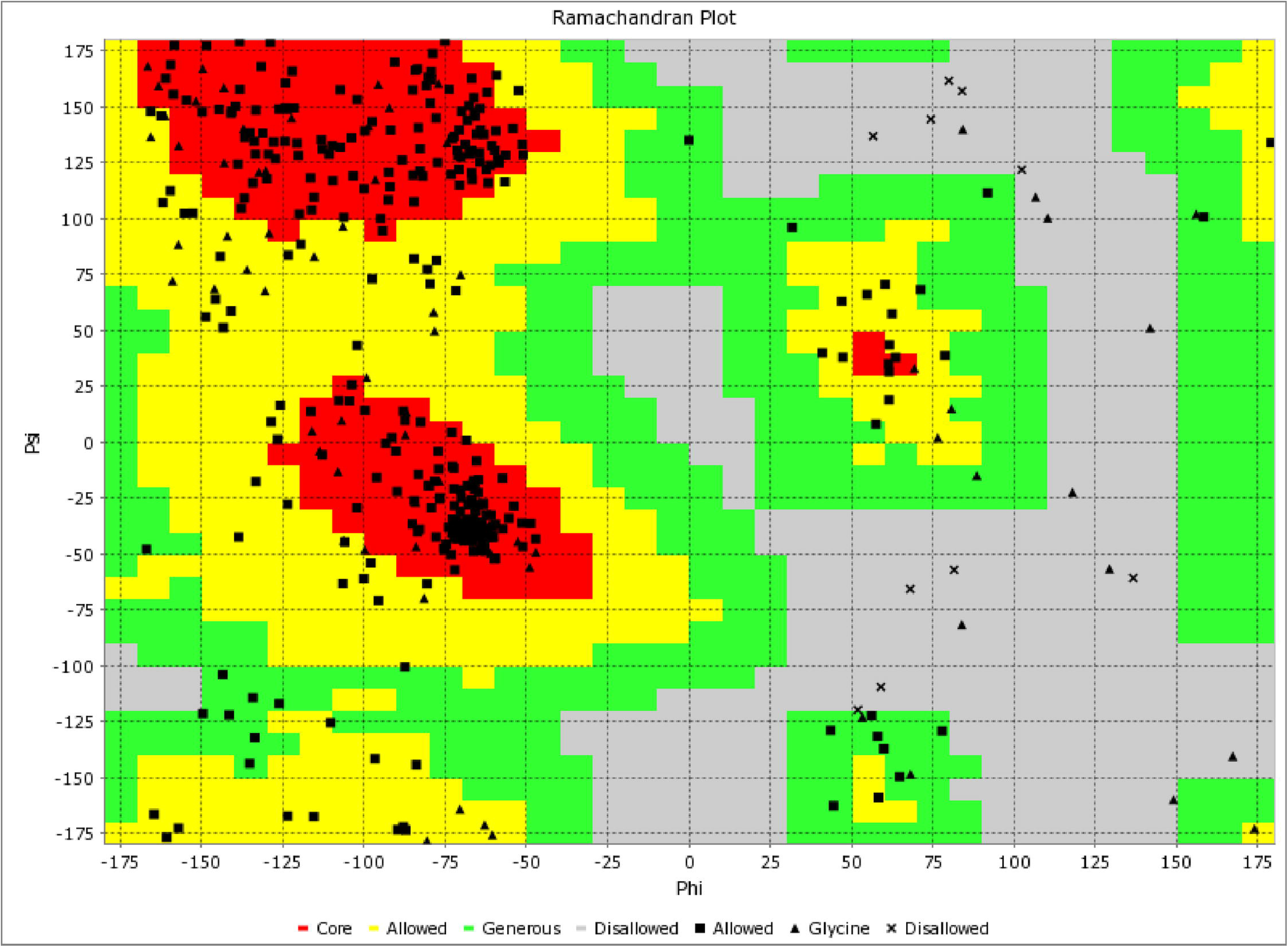
Validation: Ramachandran plot analysis of protein residues, showing 75.6 % in favoured, 21.7 % in allowed, and 2.6 % in disallowed regions

### 3.9 Predicted discontinuous B-cell epitopes

The prediction of discontinuous epitopes requires the 3D structural information of the protein or polypeptide. Thus, ElliPro server was used to predict three discontinuous B-cell epitopes for the MEV indicating its residues, number of residues, and scores (Table 8). The 3D representation of putative B-cell epitopes is shown in Fig 6 with yellow surface.

**Fig 6.**
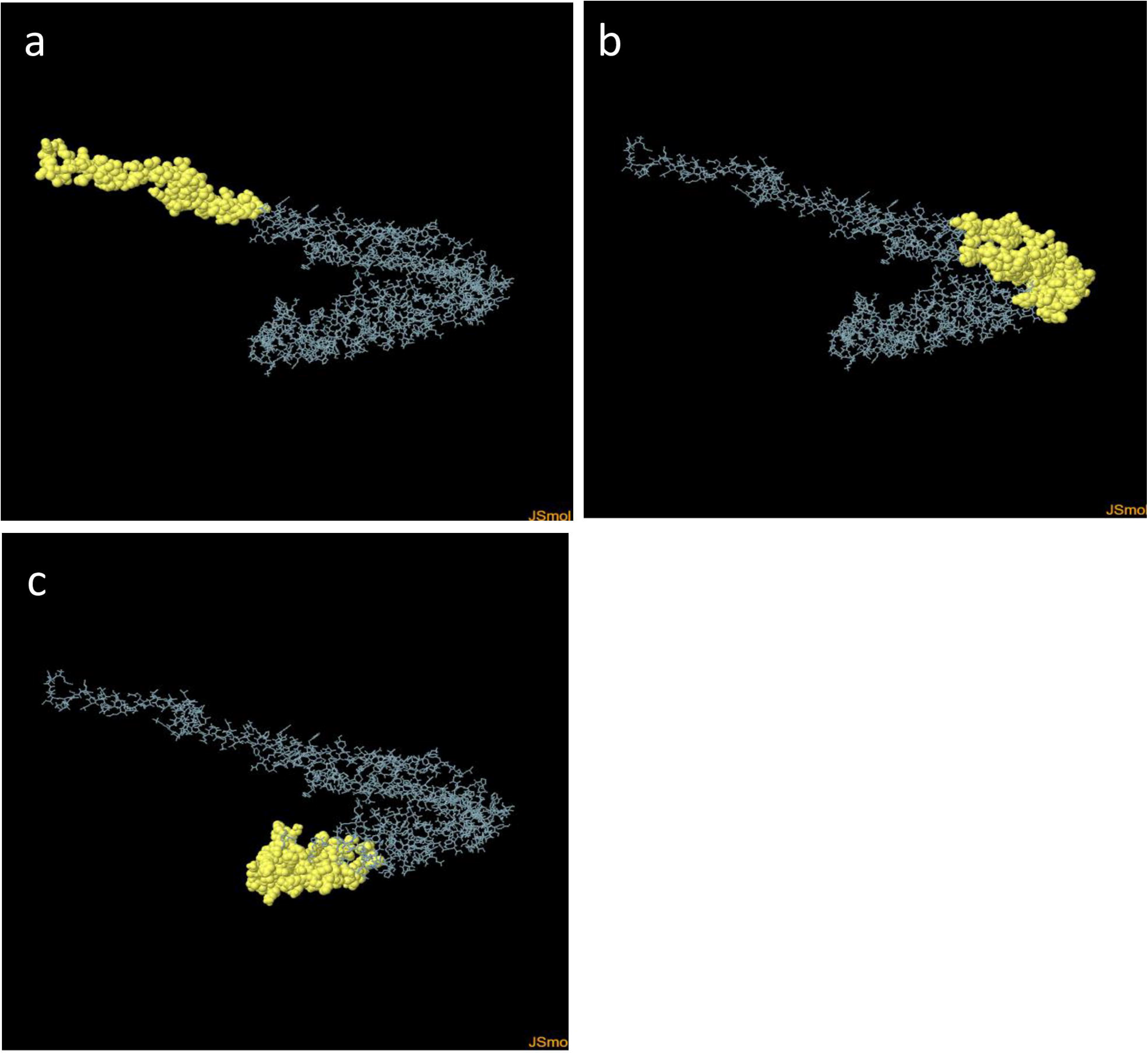
3D representation of the discontinuous B-cell epitopes of MEV. The yellow surface shows discontinuous B-cell epitopes, while the grey stick represents the bulk of the polyprotein. (a) With 61 residues and a score of 0.869, (b) with 96 residues with a score of 0.681, and (c) with 100 number of residues with a score of 0.632

**Table 7.**
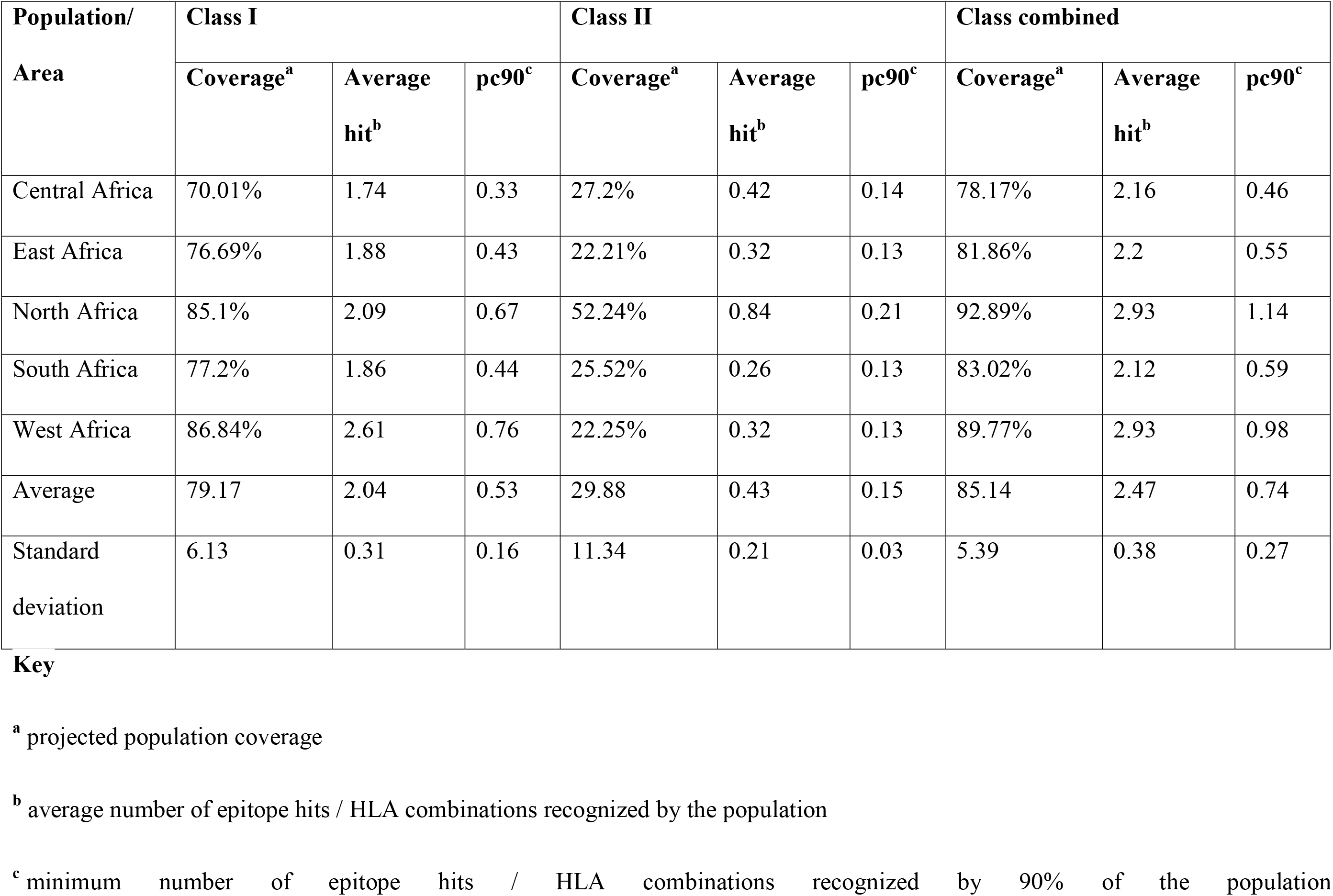
Percentage population Coverage of MHC class I and MHC class II epitopes in different regions in Africa.

**Table 8.**
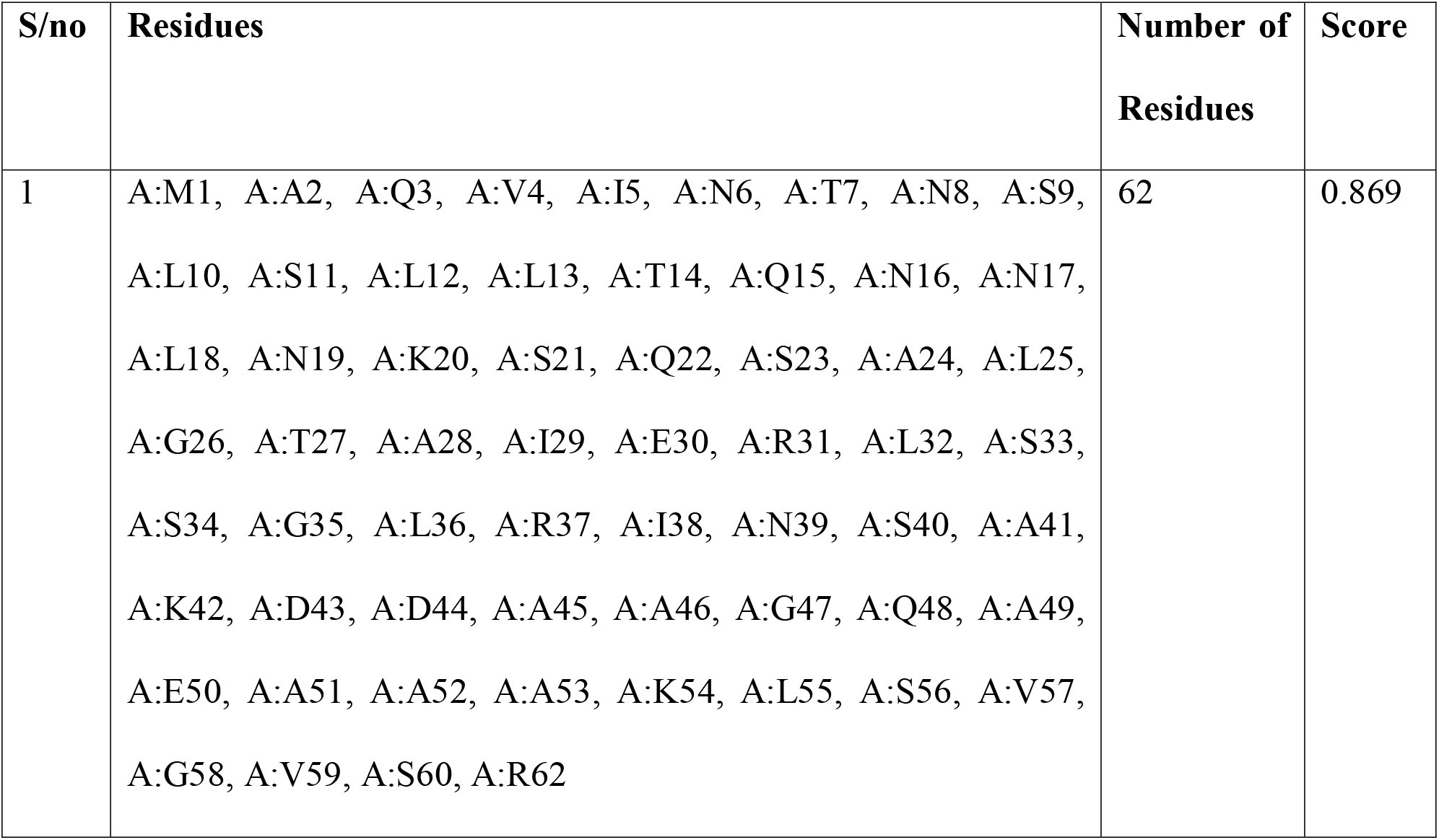
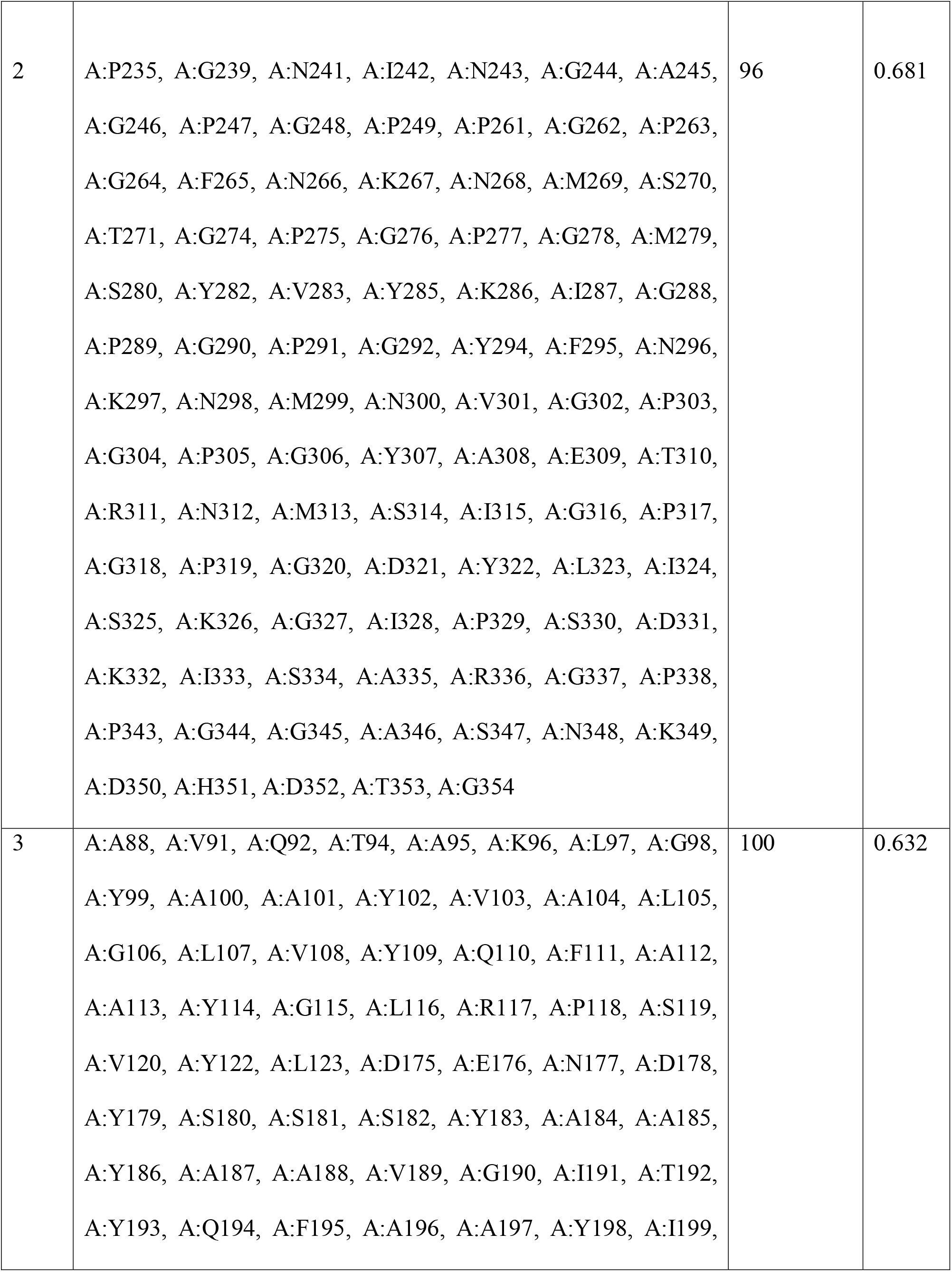
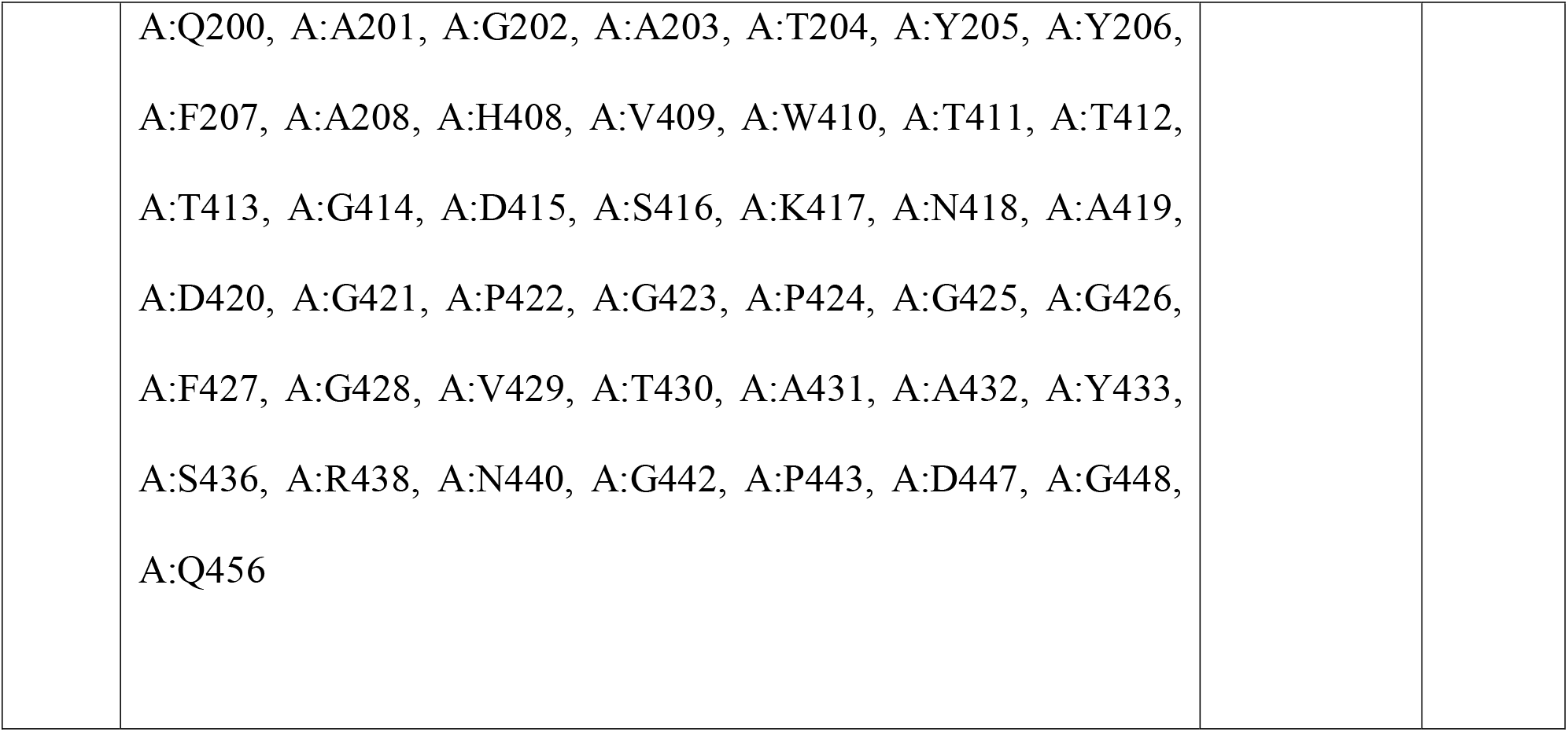
Predicted discontinuous B-cell epitopes residues of the designed MEV, predicted by ElliPro.

### 3.10 Protein-protein docking and protein-protein binding affinity simulation

Results from the ClusPro web server provided a protein-protein docking complex with the best model selected based on their lowest energy level score (Table 9). The 3D tertiary structural models of the selected toll-like receptors multi-epitope vaccine docked complexes are in Fig 7. ChimeraX software was used for visualization.

**Fig 7.**
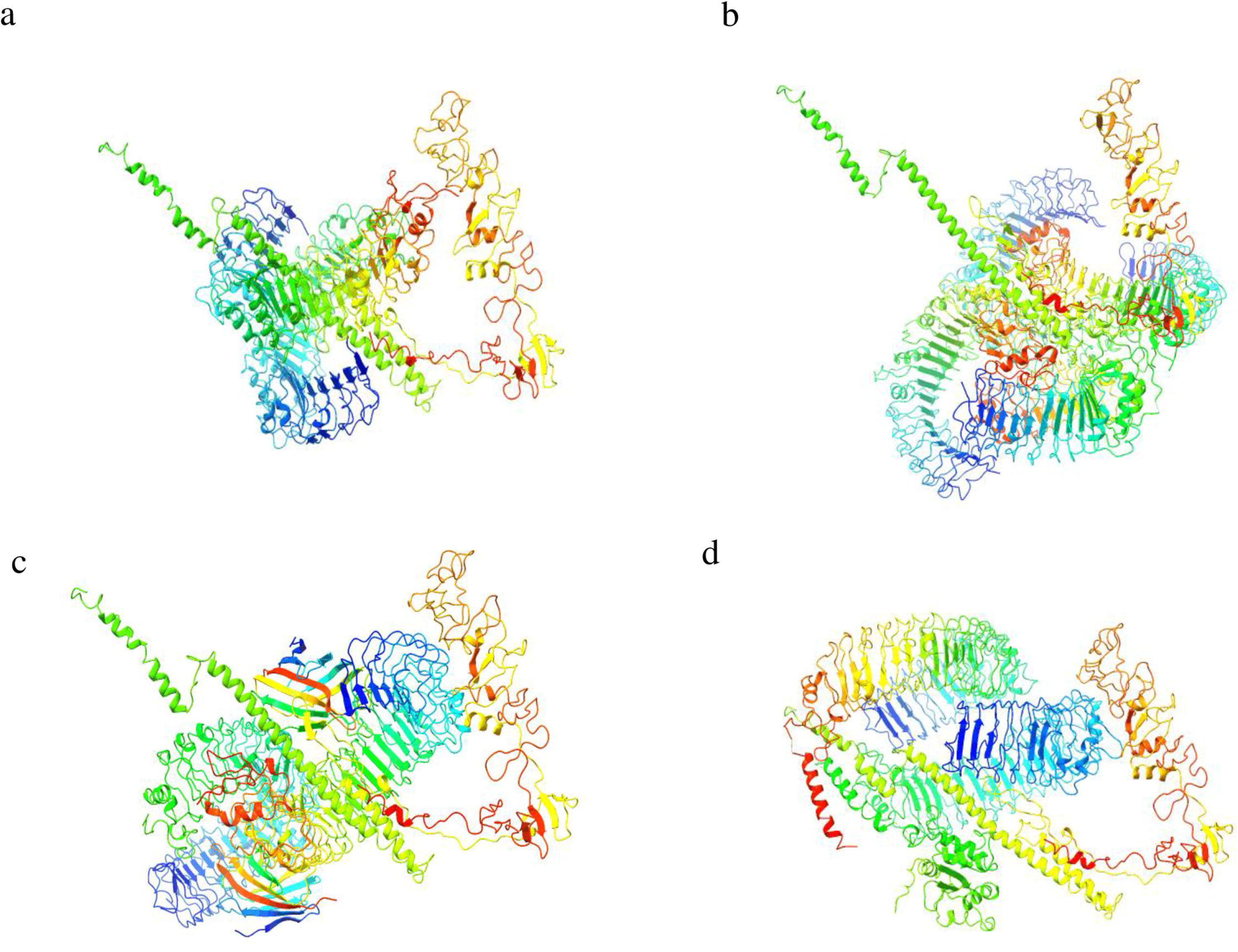
The ClusPro protein–protein docking of multi-epitope vaccine (MEV) model with Toll-like receptors (TLR). (a) MEV construct with TLR-1 (PDB Id: 6NIH), (b) MEV construct with TLR-2(PDB ID: 6NIG), (c) MEV construct with TLR-4 (PDB ID: 3FXI) and (d) MEV construct with TLR-5 (PDB ID: 3JOA). All models were visualized and coloured using ChimeraX software

**Table 9:**
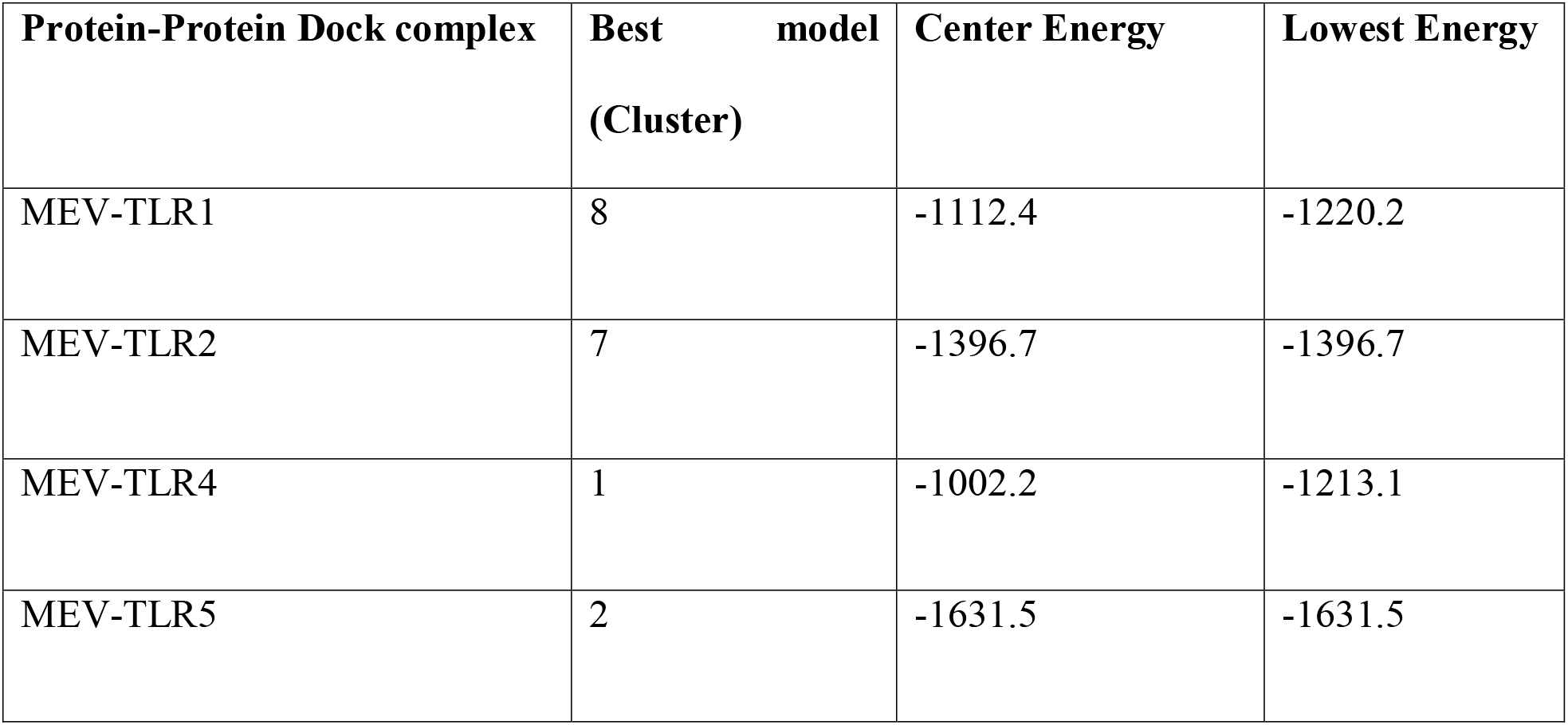
Result for protein-protein docking on the ClusPro 2.0 server.

### 3.11 Molecular dynamics simulation and protein-protein binding affinity analysis

Molecular dynamics simulations and protein-protein binding affinity analyses were conducted on the TLR-MEV docked complexes using the iMODS tool. This analysis evaluated the stability and physical movements of all TLR-MEV docked complexes, as illustrated in Figures 8 to 11. The main-chain deformability indicates the ability of a molecule to deform at each of its residues, with regions of high deformability identifying the location of the chain ‘hinges’. The B-factor values, calculated by normal mode analysis and proportional to the Root Mean Square, quantify the uncertainty of each atom in the complex.

**Figure 8.**
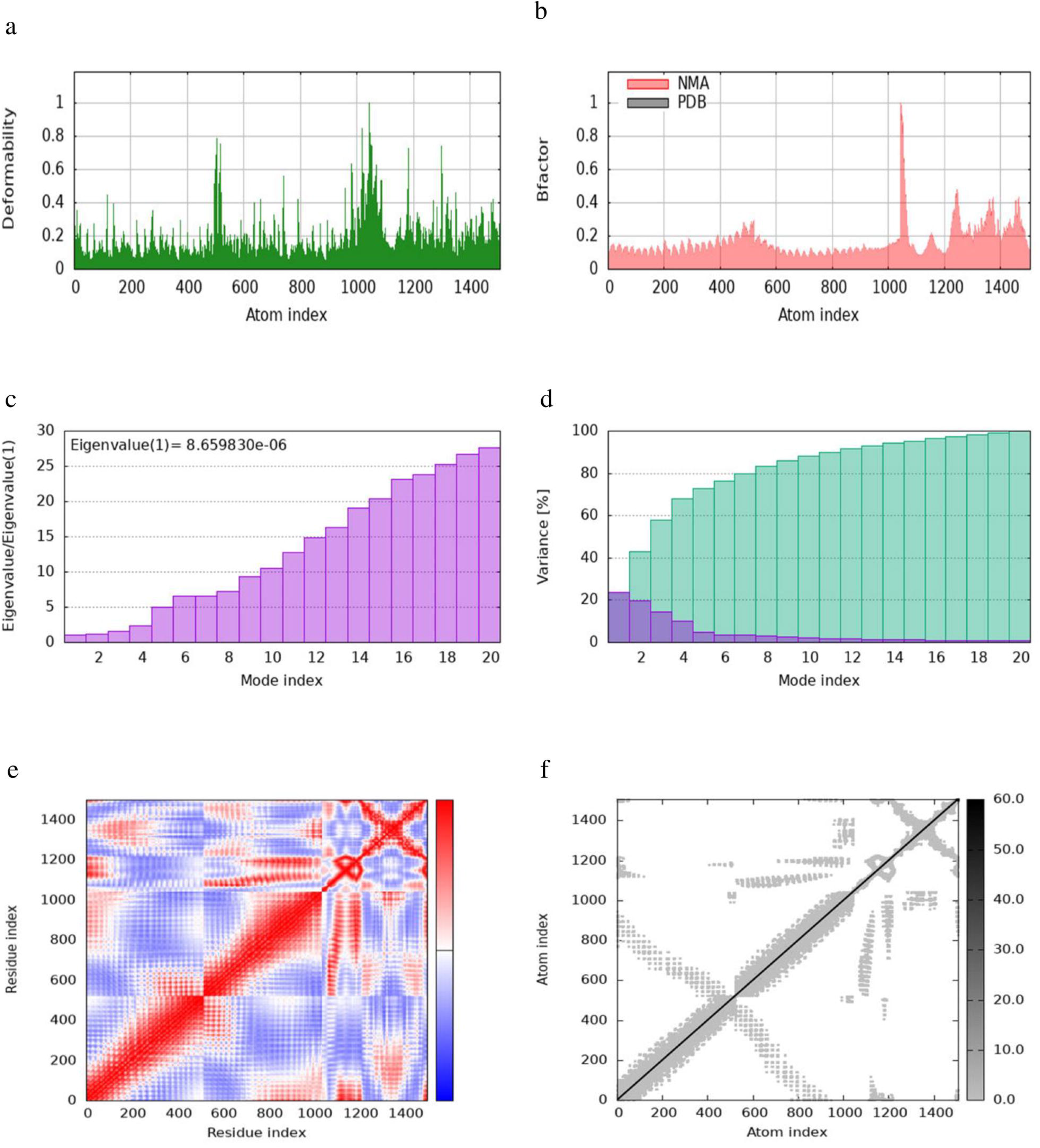
iMODS Simulation of the TLR1-MEV Docked Complex. (a) Simulation of main-chain deformability reveals the flexibility at each residue. (b) B-factor values measure the positional uncertainty of each atom in the structure. (c) Eigenvalues indicate the energy required for structural deformation. (d) The variance associated with each normal mode is inversely proportional to its eigenvalue, with individual variances shown in purple and cumulative variances in green. (e) The covariance matrix shows the degree of interaction between pairs of residues. (f) The elastic network model depicts connections between atoms using springs, with darker shades indicating stiffer springs.

**Figure 9.**
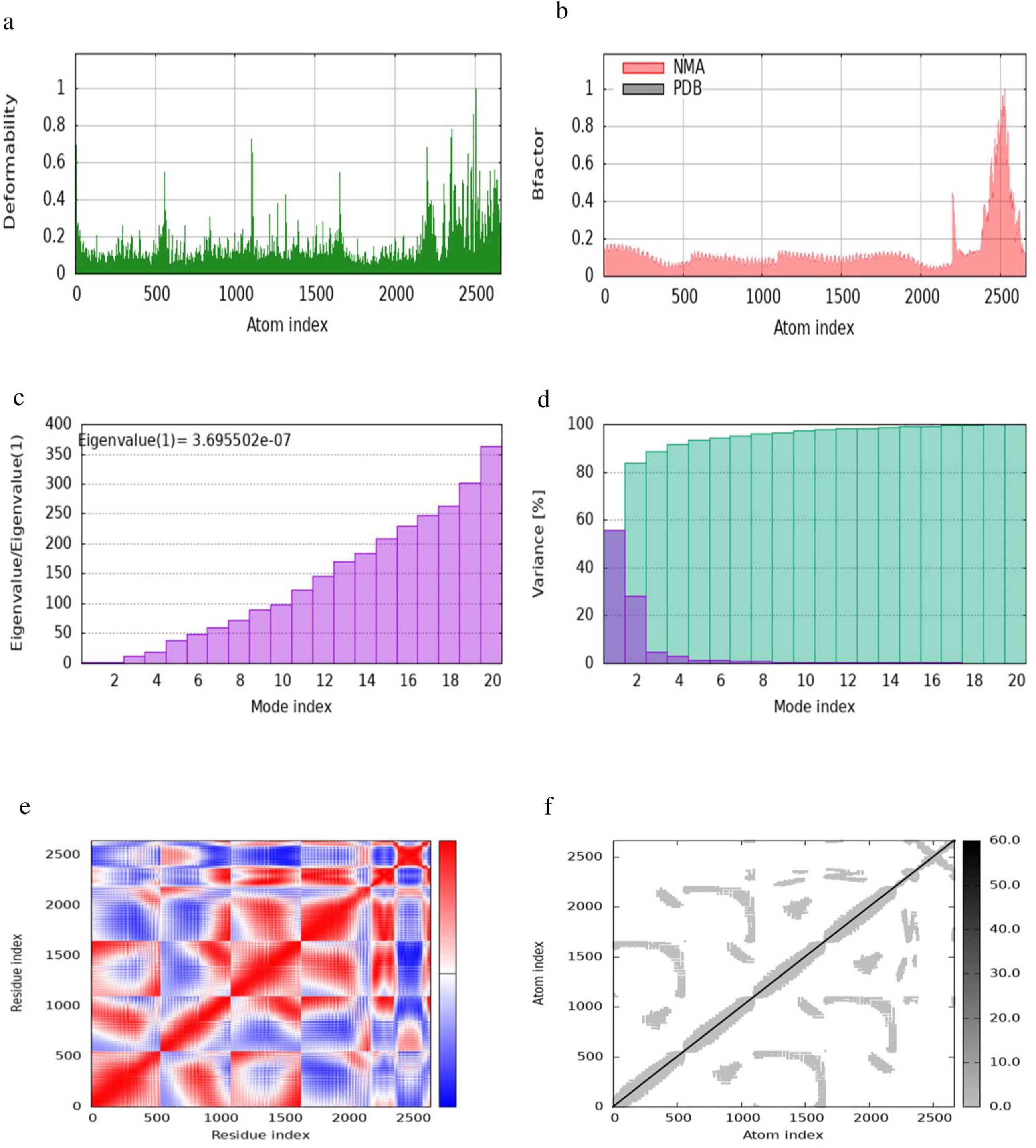
iMODs simulation of TLR2-MEV docked complex. (a) Simulation of main-chain deformability reveals the flexibility at each residue. (b) B-factor values measure the positional uncertainty of each atom in the structure. (c) Eigenvalues indicate the energy required for structural deformation. (d) The variance associated with each normal mode is inversely proportional to its eigenvalue, with individual variances shown in purple and cumulative variances in green. (e) The covariance matrix shows the degree of interaction between pairs of residues. (f) The elastic network model depicts connections between atoms using springs, with darker shades indicating stiffer springs.

**Figure 10.**
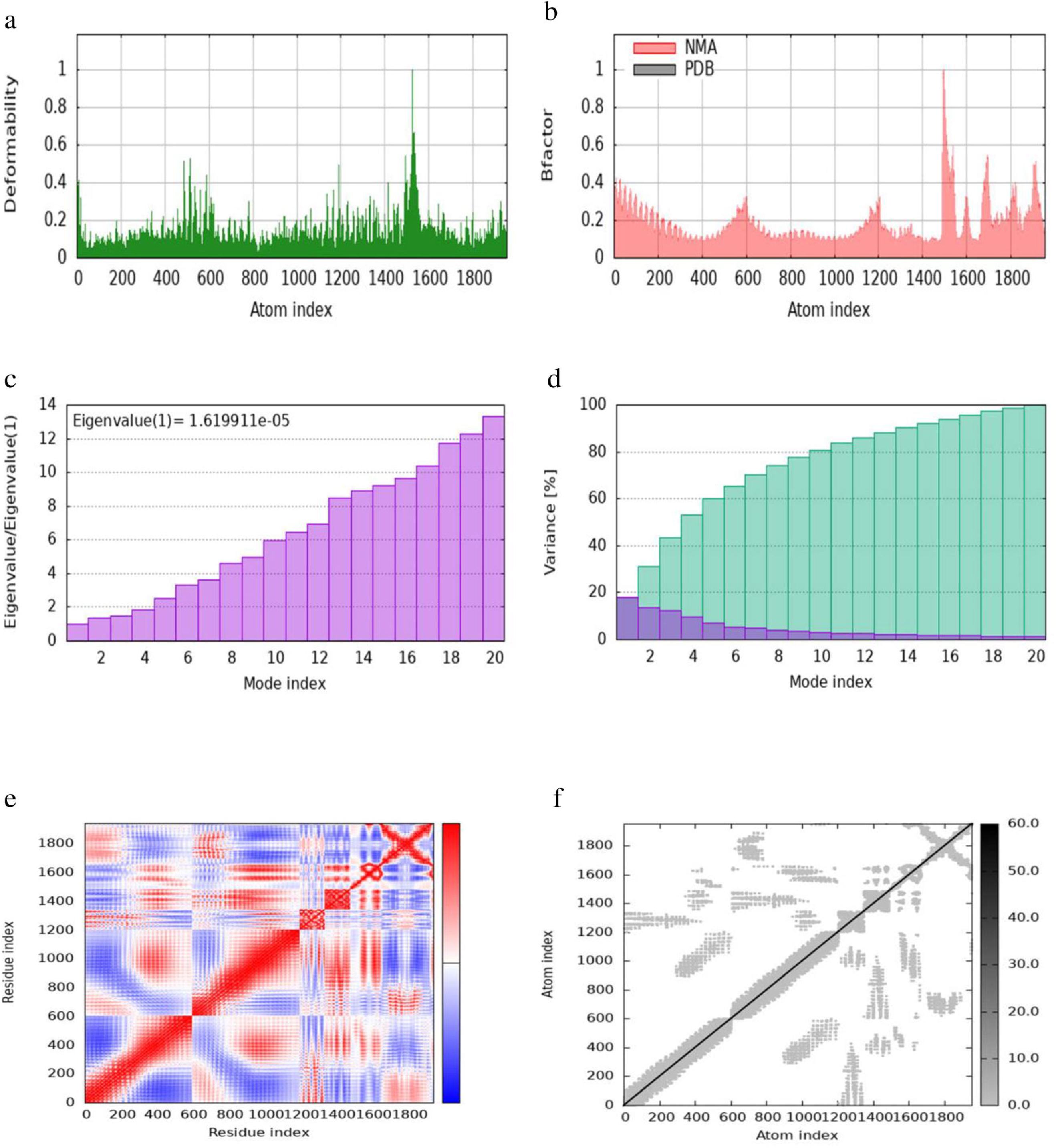
iMODs simulation of TLR4-MEV docked complex. (a) Simulation of main-chain deformability reveals the flexibility at each residue. (b) B-factor values measure the positional uncertainty of each atom in the structure. (c) Eigenvalues indicate the energy required for structural deformation. (d) The variance associated with each normal mode is inversely proportional to its eigenvalue, with individual variances shown in purple and cumulative variances in green. (e) The covariance matrix shows the degree of interaction between pairs of residues. (f) The elastic network model depicts connections between atoms using springs, with darker shades indicating stiffer springs.

**Figure 11.**
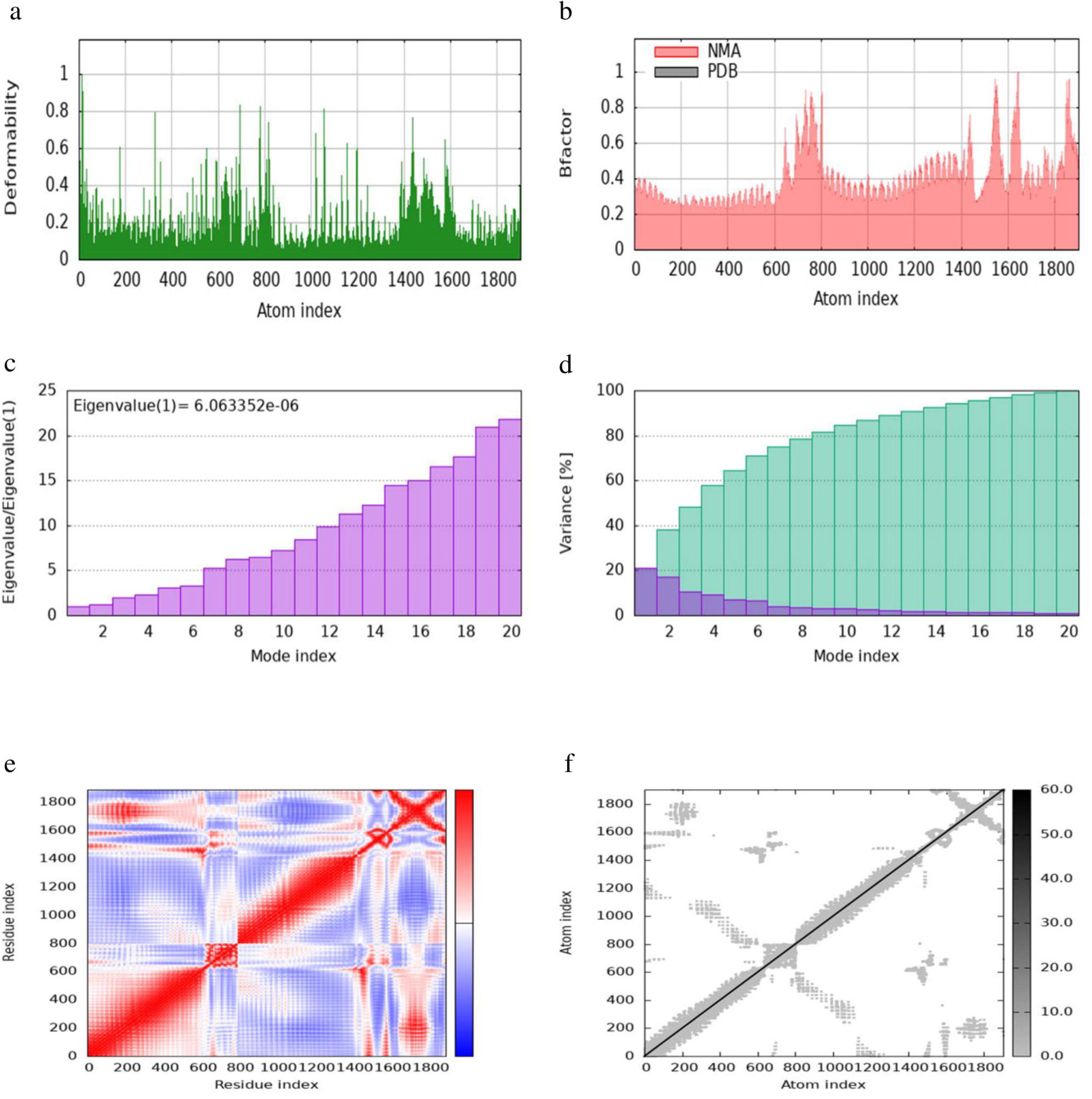
iMODs simulation of TLR5-MEV docked complex. (a) Simulation of main-chain deformability reveals the flexibility at each residue. (b) B-factor values measure the positional uncertainty of each atom in the structure. (c) Eigenvalues indicate the energy required for structural deformation. (d) The variance associated with each normal mode is inversely proportional to its eigenvalue, with individual variances shown in purple and cumulative variances in green. (e) The covariance matrix shows the degree of interaction between pairs of residues. (f) The elastic network model depicts connections between atoms using springs, with darker shades indicating stiffer springs.

The eigenvalue of the docked complex represents the energy required to deform the structure, with the following values obtained: 8.6598 × 10 for TLR1-MEV, 3.6955 × 10 for TLR2-MEV, 1.6199 × 10 for TLR4-MEV, and 6.0634 × 10 for TLR-MEV. The covariance matrix plot shows the correlations between pairs of residues, with red indicating correlated, white indicating uncorrelated, and blue indicating anti-correlated pairs. The elastic network model plot illustrates the connections between atoms, where darker shades of grey represent more rigid connections. For the binding affinity analysis, the Prodigy bioinformatics tool was used, revealing a Pearson’s Correlation coefficient (r) of 0.73 (p-value < 0.0001) between the predicted and experimental values, and a Root Mean Square Error (RMSE) of 1.89 kcal mol ¹. Intermolecular contacts were defined as residues with heavy atoms within 5.5 L of each other. The Prodigy server predicted 54 intermolecular contacts for the MEV-TLR1 complex, 101 for MEV-TLR2, 168 for MEV-TLR4, and 50 for MEV-TLR5. The binding affinity score (ΔG in kcal mol¹) was negative for all docked complexes, as shown in Table 10.

**Table 10.**
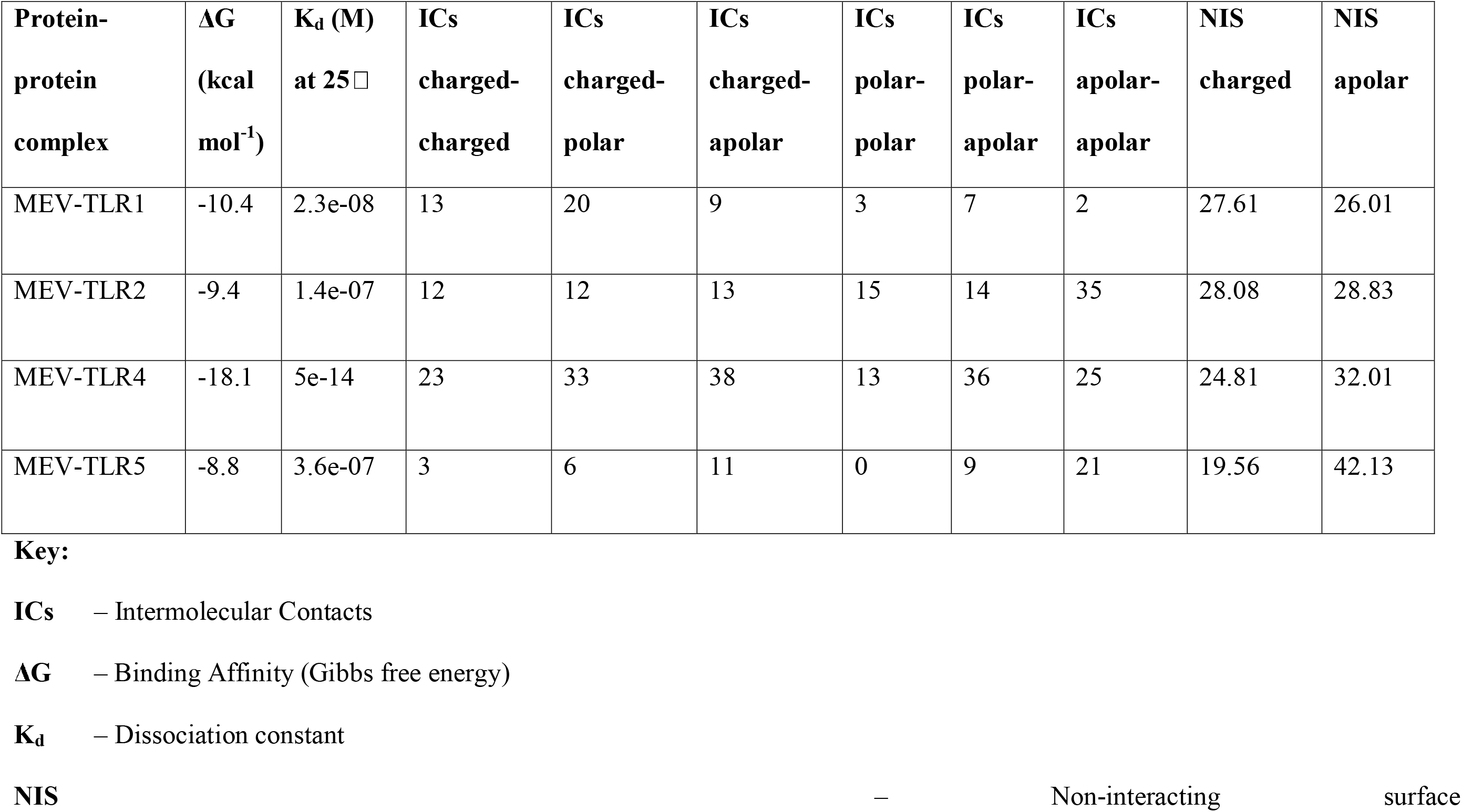
Prediction of binding Affinity and Dissociation constant.

### 3.12 Immune Simulation

The administration and repeated exposure to the MEV vaccine led to a significant increase in the antibody response, coupled with a gradual decline in antigen levels over time. A notable IgM and IgG humoral response indicated some levels of seroconversion. The humoral responses for both IgM and IgG were more pronounced after booster doses compared to the initial dose, with IgM levels being higher than IgG (and its isotypes) (Fig. 12a). Following the initial dose, both CD8+ T-cell and CD4+ responses were actively stimulated. CD4+ T-helper lymphocytes peaked after the second booster dose, with a significant increase in the active entity state and Th-memory phenotype observed after the first booster, remaining stable with minimal decay over 100 days (Fig. 13a and Fig. 13b). Similarly, the prime dose induced a significant spike in the IFN-g response (associated with both CD8+ T-cell and CD4+ Th1 response) and a substantial increase in IL-2, IL-10, IL-12, and TGF-b (Transforming Growth Factor-beta) cytokine responses, linked to the T-reg phenotype (TGF-b and IL-10). These responses persisted after the two booster doses, with IFN-g response maintained and IL-2, IL-10, IL-12, and TGF-b levels rising significantly (Fig 15b). These immune readouts are typically associated with a broader peak in antigen quantification.

**Fig 12.**
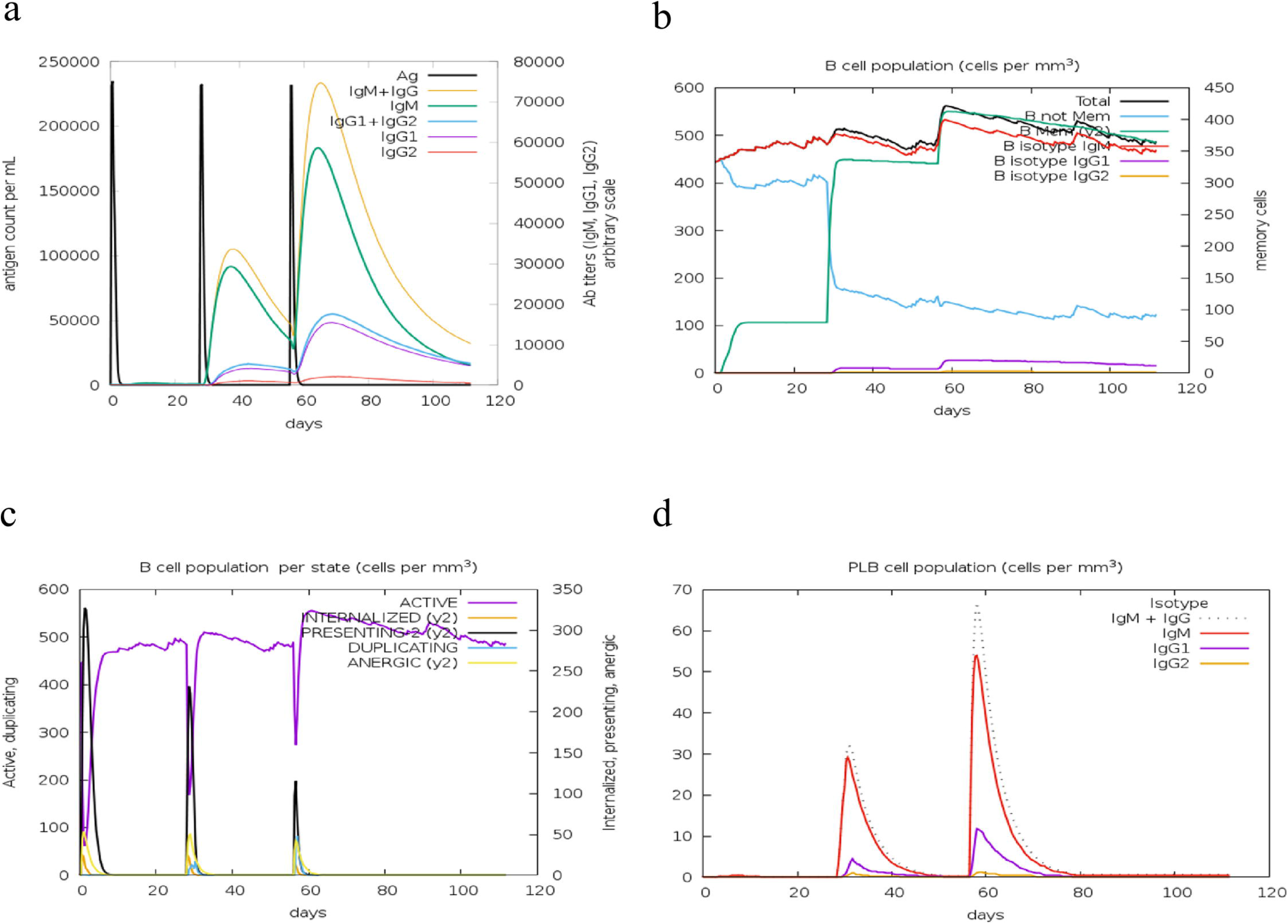
Distribution of B-lymphocytes and their isotypes. (a) Antigen and immunoglobulin levels, with antibodies classified by isotype. (b) Total B-lymphocyte count, memory cells, and isotypes IgM, IgG1, and IgG. (c) B-lymphocyte population by entity-state, showing counts for active, class-II presenting, antigen-internalized, duplicating, and anergic states. (d) Plasma B lymphocyte (PLB) count divided by isotype (IgM, IgG1, and IgG2).

**Fig 13.**
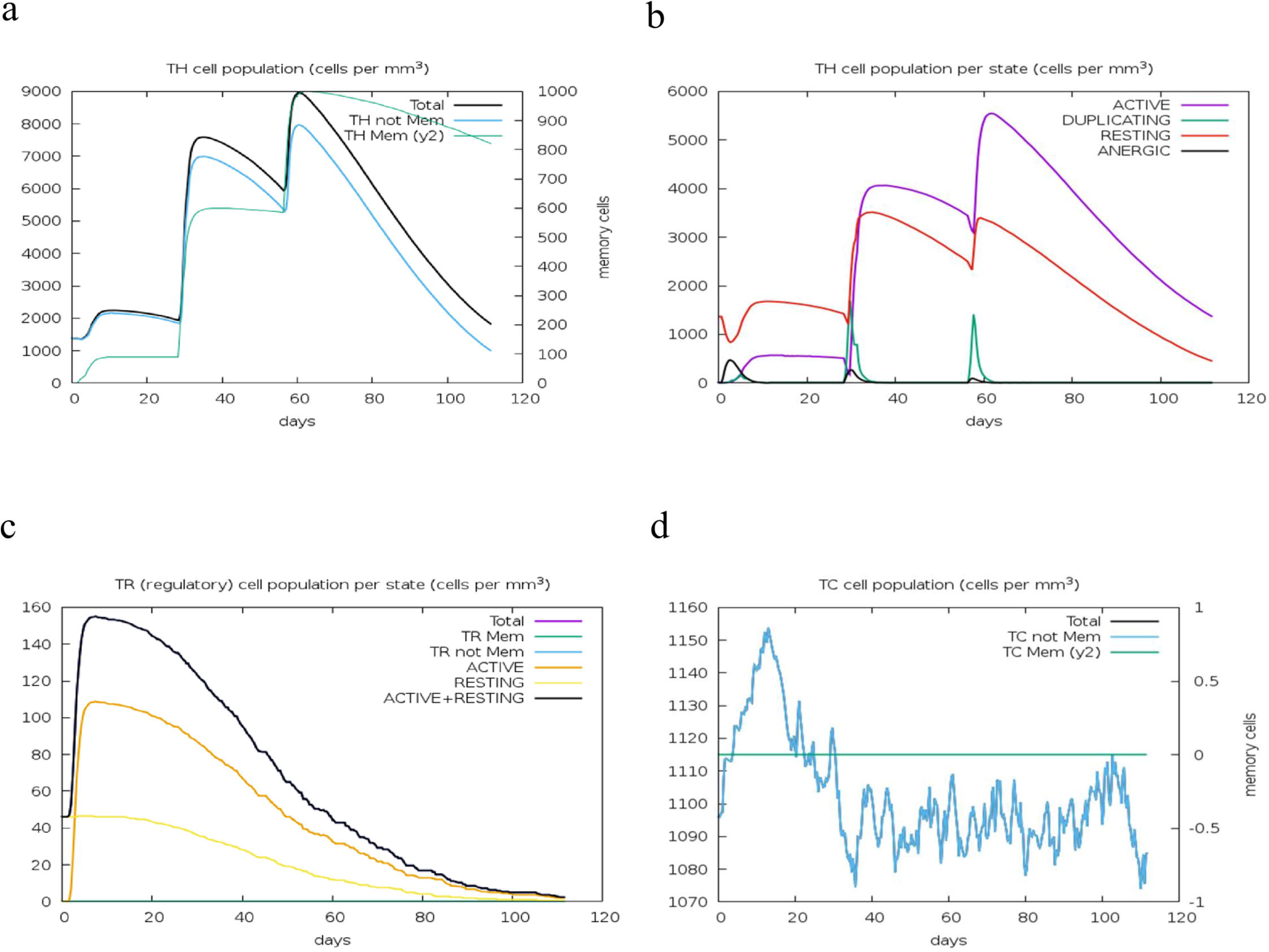
T-lymphocyte populations. (a) CD4+ T-helper lymphocyte counts, showing total and memory cells. (b) CD4+ T-helper lymphocyte counts by entity state, including active, resting, anergic, and duplicating states. (c) CD4+ T-regulatory lymphocyte counts, with total, memory, and entity-state counts plotted. (d) CD8+ T-cytotoxic lymphocyte counts, showing total and memory cells.

Analysis of the B-cell population per cell showed overall B-cell population and B-cell memory responses were higher and stable showing minimal decay. B-cell memory was effectively elicited, peaking after the second booster dose and remaining stable over 100 days. Similarly, the active B-cell population was higher and stable over the same period (Fig. 12b and Fig. 12c). There was also an increase in the activity of dendritic cells, natural killer cells, and macrophages throughout the simulation (Fig. 14b, Fig. 14c, and Fig. 14d). This indicates the vaccine’s effectiveness in stimulating the appropriate immunological components for an effective response. Overall, these results demonstrated a steady and concomitant increase in immune responses over time with each vaccination schedule.

**Fig 14.**
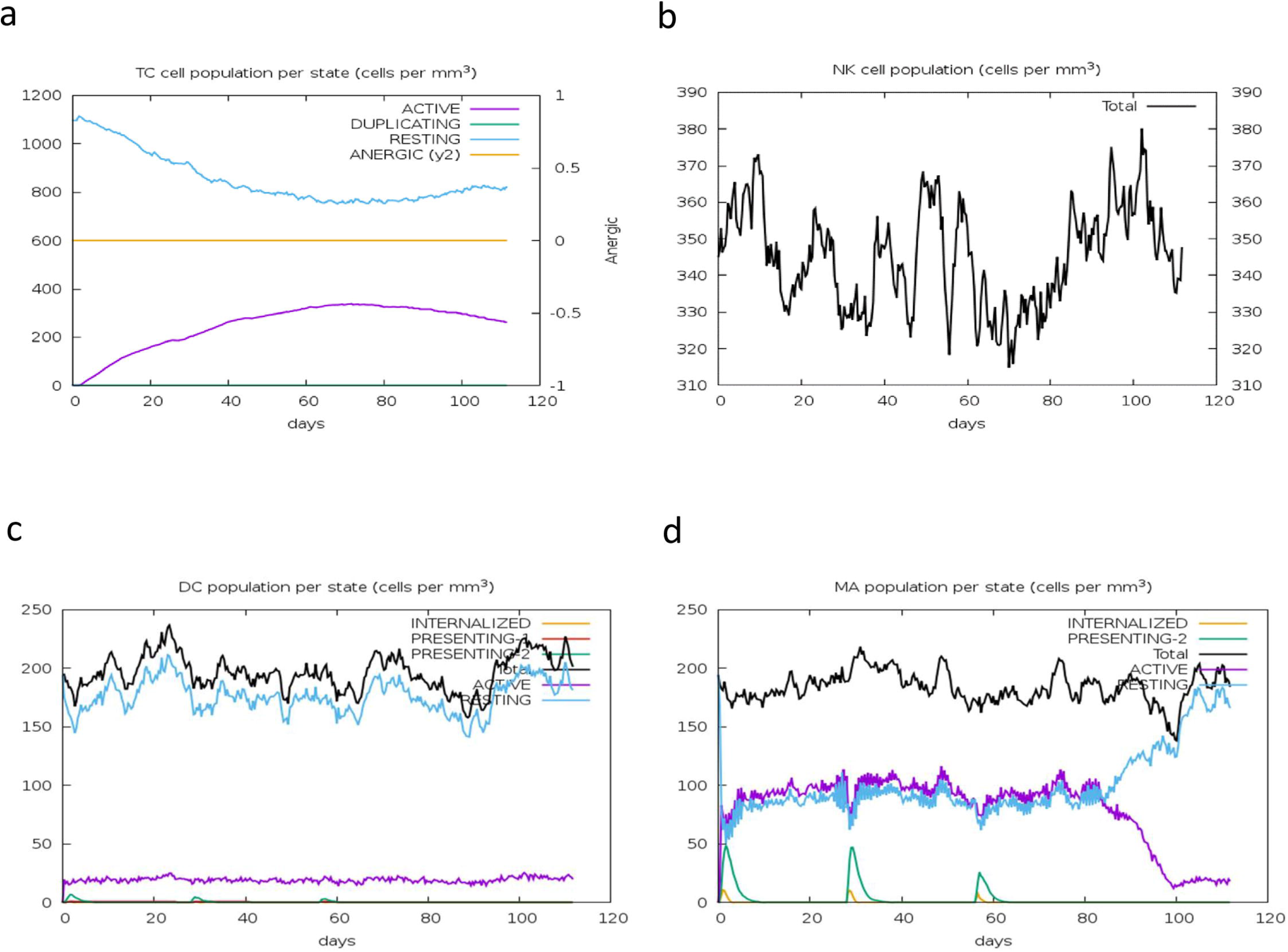
Immune cell activity. (a) CD8+ T-cytotoxic lymphocyte counts by entity-state. (b) Total count of natural killer cells. (c) Dendritic cells, which present antigenic peptides on both MHC class-I and class-II molecules, with the total number broken down into active, resting, internalized, and antigen-presenting states. (d) Macrophage counts, including total, internalized, class-II presenting, active, and resting states.

### 3.13 Codon adaptation and *In-Silico* plasmid cloning

The optimized vaccine codon sequence was 1,386 nucleotides long. The initial GC content was 40.89%, which increased to 70.78% after optimization using the Java Codon Adaptation Tool (JCAT), enhancing translational efficiency. The calculated codon adaptation index was 0.956. The NEBcutter Version 3.0.17 software was used to identify restriction enzyme sites on the JCAT-optimized MEV sequence. The MEV coding DNA was inserted between HindIII and BamHI in the multiple cloning site (MCS). *In-silico* cloning was performed using SnapGene software, resulting in a Plasmid-MEV clone of 6844 bp (Fig. 16).

The red segment represents the coding gene for the multi-epitope, while the black segment indicates the plasmid vector backbone. The plasmid DNA (pDNA) size is 6844 bp.

## 4.0 Discussion

Outer membrane proteins (OMPs) play crucial roles in antibiotic resistance by acting as gatekeepers that regulate the influx of external molecules. These proteins are also pivotal in cellular communication and have promising applications as biosensors. Their multi-functionality makes them excellent candidates for vaccine development. This study leverages the immunogenic properties of OMPs, along with the *fliC* protein, to design an innovative multiepitope vaccine (MEV). The computational approach to designing chimeric membrane proteins aims to optimize the advantages of these molecular gateways and signaling networks [65, 66, 67].

This study identified a total of 29 epitopes with the potential to stimulate both cellular and humoral immune responses, activating T-cell and B-cell responses. Across the three classes of OMPs, 15 CTL epitopes were predicted using specified thresholds. Notably, the epitope “IEYAITPEI” was identified in both MHC class I and MHC class II epitopes of OmpA, as well as in MHC class I epitopes of OmpC. All epitopes were highly conserved with a minimum and maximum conservancy identity at 100 % (Table 4), ensuring the vaccine is efficacious against the diverse bacterial variants. All the CTL peptide epitopes also showed highly exposed residue, therefore capable of triggering a robust cytotoxic response. Typically, CD8+ T-cells recognize pathogen antigens bound to MHC-I molecules, triggering cytotoxic responses against the pathogen [68]. All selected epitopes were found to be non-toxic and non-allergenic, reducing the likelihood of the vaccine candidates causing type II hypersensitivity reactions or cellular apoptosis due to toxic reactions. Using specified thresholds and immunogenicity scores, eight HTL epitopes were predicted across the three classes of OMPs. These epitopes were further evaluated for their capacity to induce IFN-γ, as the release of IFN-γ by CD4+ helper T-cells is essential for activating effector cells and generating antibody-mediated responses to infections [31]. IFN-γ plays a crucial role in pathogen recognition and elimination, serving as a key effector in cell-mediated immunity and coordinating a plethora of antimicrobial functions. It is especially important in mediating intestinal immunity against *Salmonella* spp. [31, 69]. The study’s findings predicted that two HTL epitopes, “LAPDRRVEI” and “IEYAITPEI,” could induce the production of IFN-γ (Table 6). This finding is satisfactory as IFN-γ is expected to trigger a cascade of receptor-ligand interactions involving other cytokines and pattern recognition receptors (PRRs) such as interleukin-4 (IL-4), TNF-α, and lipopolysaccharides [31, 70]. More so, excessive levels of IFN-γ have been linked to significant tissue damage, necrosis, and inflammation, potentially exacerbating disease pathology [31].

In this study, seven linear B-cell epitopes were identified, with their positions, lengths, sequences, and antigenicity provided in Table 5. These epitopes were selected utilizing an artificial neural network (ANN) and the Kolaskar and Tongaonkar antigenicity scale method from the IEDB server, which boasts a 75% accuracy in predicting antigenic determinants of proteins. The identified B-cell epitopes are non-toxic and non-allergenic, making them suitable candidates for vaccine development. The results suggest that hydrophobic residues such as cysteine, leucine, and valine on protein surfaces are likely to possess antigenic properties [71].

When designing a vaccine, it is critical to consider the population coverage capacities of the predicted MHC class I and MHC class II T-cell epitopes. Given the presence of over a thousand human MHC alleles worldwide, it is essential to ensure that the vaccine can elicit an immune response in individuals with diverse MHC alleles [72, 73]. Our vaccine demonstrated a global population coverage of 94.91%, indicating that approximately 95 % of the global population would likely be protected if vaccinated with our construct containing the identified CD8+ and CD4+ T-cell epitopes. Europe showed the highest coverage at 97.52% (Fig 1), suggesting significantly higher protection from our vaccine chimera in this region compared to others. Given that the *Salmonella* Kentucky *ST198* MDR clone was first identified in Egypt, and is now widespread in Africa [3, 74, 75, 76], we were particularly interested in the vaccine’s performance across different African regions. The vaccine coverage in Africa was substantial at 85.14%, with CD8+ T-cell epitopes achieving higher coverage than CD4+ T-cell epitopes (Table 7). North Africa exhibited the highest coverage on the continent, with a combined average of 92.89% for both classes. The CD4+ T-cell coverage in North Africa was notably higher at 52.24%, nearly double that of other African regions for this T-cell class (Table 7). The variations in regional coverage may be attributed to differences in ethnicity and race. North Africa’s mixed population, including Arabs, Caucasoid, Blacks, and the Tunisian Berbers, shares more allele similarities with Europe and Africa combined than with the rest of Black Africa. Previous studies have suggested that such variations in population coverage could be due to the IEDB server population coverage tool having fewer allele entries from continents like Africa and Asia compared to Europe and America [37]. This discrepancy might also explain the variations observed in our study.

The multiepitope vaccine was constructed by integrating predicted T-cell epitopes, B-cell epitopes, and the *fliC* protein, interconnected using AAY, GPGPG, and EAAK protein linkers respectively (Fig 2 and Fig 4). These linkers are crucial for preserving the immunogenicity and properties of proteins within the MEV chimera. The absence of linkers in MEV construction could lead to the formation of a novel protein with unknown characteristics or the generation of neo-epitopes and junctional epitopes [77, 78]. From an immunological perspective, protein fusions are advantageous as each antigen in the fusion can benefit from additional T-cell help provided by partner T-cell epitopes, resulting in enhanced antigen-specific immunogenicity and antibody titers [79]. The physicochemical analysis of the MEV construct indicated a highly antigenic, non-allergenic, and non-toxic chimera with a molecular weight of 47.343 kD and a theoretical pI of 9.150. The molecular weight of peptides used in MEV should ideally be less than 110 kD, and the theoretical pI score suggests the basic nature of our MEV chimera [80, 81]. The MEV was predicted to be thermo-stable, soluble (hydrophilic), and predominantly composed of random coils (69.91%) in its protein secondary structure (Fig 3). The abundance of random coils in the MEV underscores its potential for efficient antigenicity and may also reflect the natural antigenic nature of the native unfolded alpha-helices secondary structure [18, 81, 82]. Analysis through the Ramachandran plot demonstrated that over 95% of residues in the MEV model were located within the allowed region (Fig 5), indicating an overall satisfactory 3D model. Verification through ERRAT further confirmed the stereo-chemical accuracy of the MEV’s 3D structure.

The evaluation of protein-protein interactions demonstrated that the least energy values of all our docked complexes indicate a stable TLR-MEV interaction (see Table 9). Molecular simulation analysis further suggested a low probability of complex deformation during immune response activity, as evidenced by their maximum eigenvalues (refer to Figures 8 to 11) [58]. To assess the likelihood of these complexes forming, we calculated the Gibbs free energy (ΔG). In thermodynamic terms, Gibbs free energy (ΔG) or the binding affinity of a complex provides a quantitative measure for determining actual interactions within a cell under specific conditions [83]. The study results revealed that all TLR-MEV docked complexes were energetically favorable, indicated by the negative Gibbs free energy (ΔG) values and multiple intermolecular contacts (see Table 10). This confirms that our MEV can effectively induce, bind, and sustain adequate immunological interactions.

The C-ImmSim server was utilized to predict the immune response stimulated by our MEV. The simulation results closely align with observed real-world immunological responses [18]. B-cell lymphocytes are known to play a crucial role in antibody production and the generation of protective immunity [84]. Upon administration and repeated exposure to the MEV, there was a significant elicitation of B-cell isotypes and B-cell memory cells, which endured for an extended period (see Figure 12a). Furthermore, the MEV administration triggered an adequate response from CD4+ T-helper cells and CD8+ T-cells. T-cell subsets are vital components of adaptive immunity in *Salmonella* infection, crucial for clearing primary infection and resisting subsequent challenges [84]. Given Salmonella’s intracellular nature, recent research underscores the collaboration between CD4+ helper cells and B-cell responses in combating *Salmonella* infection [85], as demonstrated by our MEV simulations. There was a concurrent production of NK cells, dendritic cells, and macrophages following the initial dose of the MEV. Significant levels of cytokines, including IFN-γ, TGF-β, IL-2, IL-10, and IL-12, were elicited and maintained (see Figure 15b). The overall activity of our MEV suggests that the selected epitopes effectively stimulated the necessary immunological responses for *Salmonella* protection and clearance post-infection.

**Fig 15.**
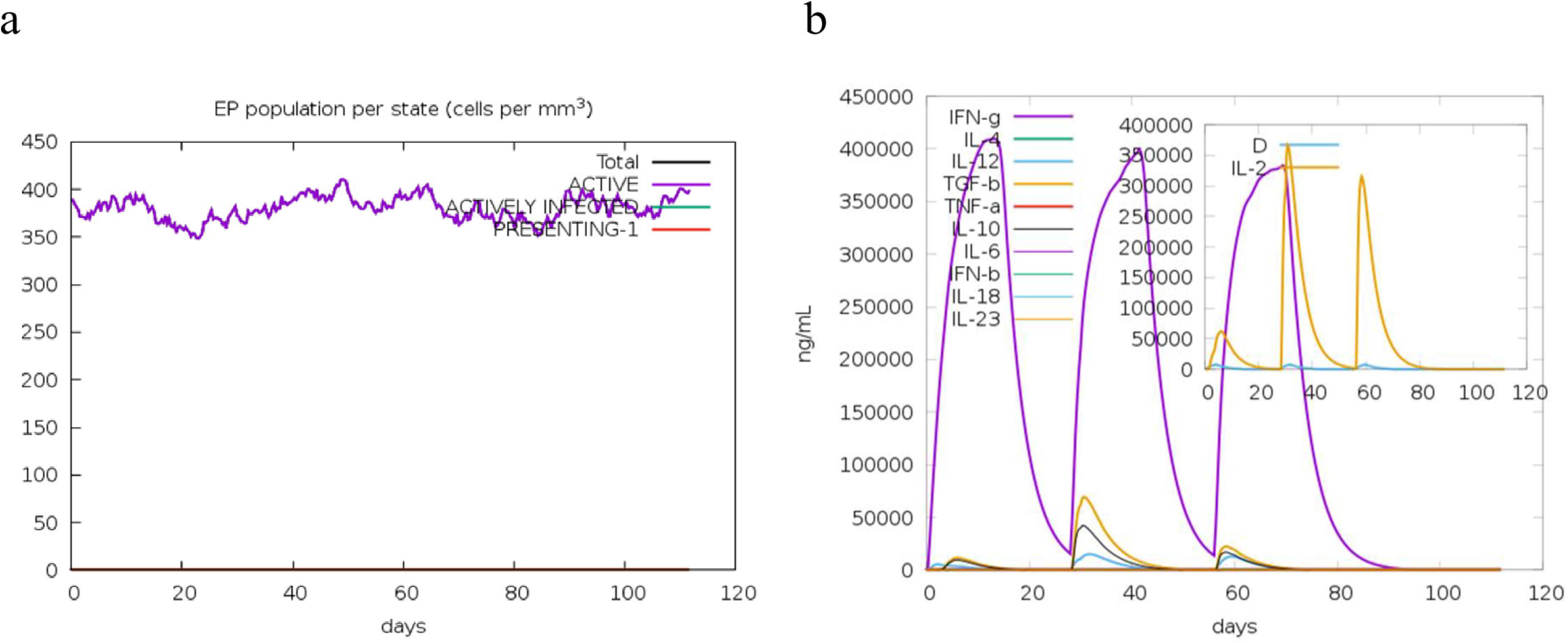
Epithelial cells and cytokines. (a) Epithelial cell counts, broken down into active, antigen-infected, and class-I MHC presenting states. (b) The concentration of cytokines and interleukins, with the inset plot showing danger signal (D) and leukocyte growth factor IL-2.

**Fig 16.**
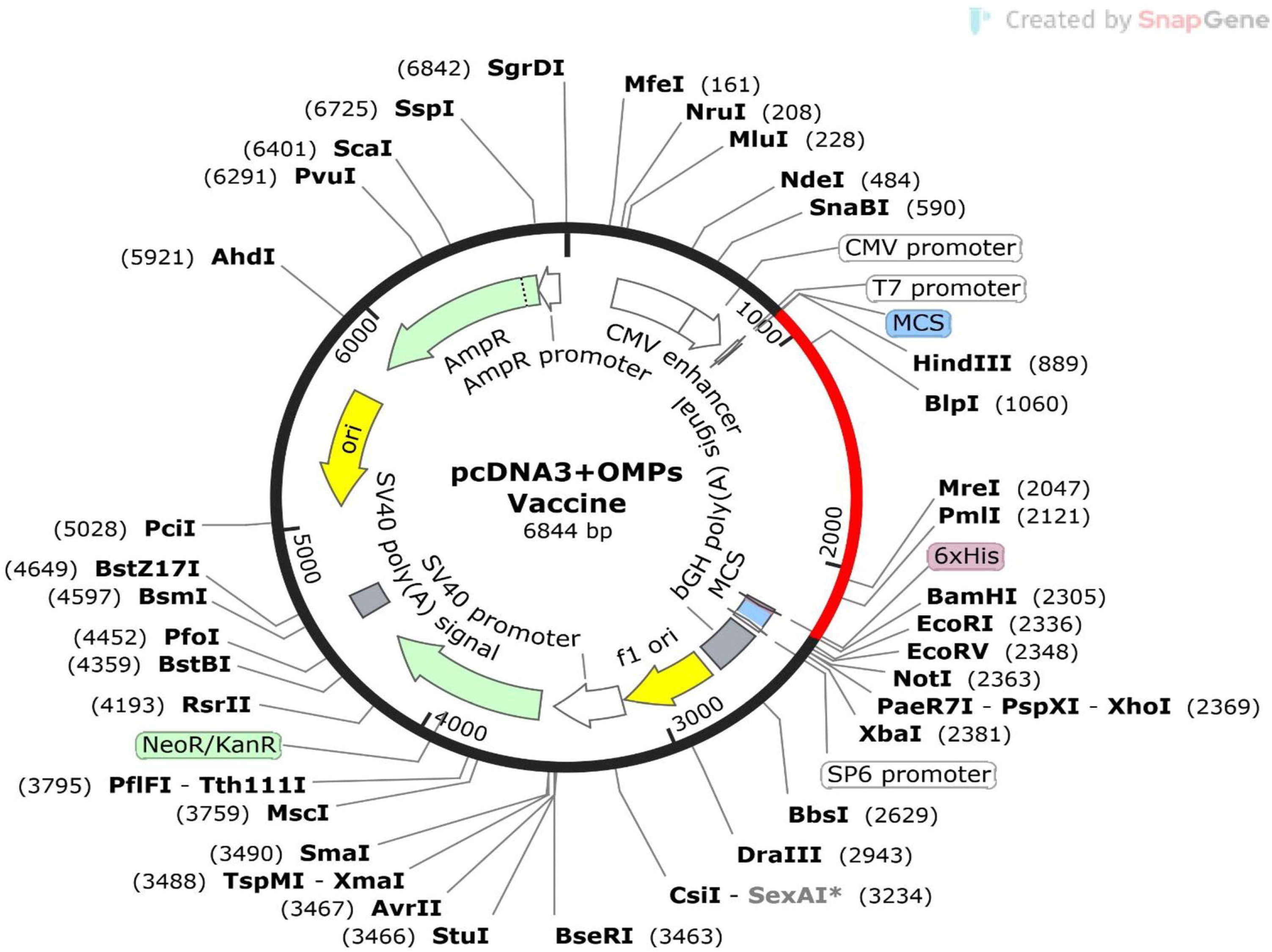
*In-Silico* Cloning of the Final MEV in pcDNA3.1_CT-GFP Expression Vector.

In mammalian expression systems, codon optimization involves increasing the proportion of preferred mammalian codons in target genes. The notably high GC content (70.78%) of the enhanced MEV protein sequence and a CAI score exceeding 0.95 indicate the likely efficient expression of our target genes in humans. Codon usage in humans is generally higher than in most non-mammalian species, with more than 95% GC3 content. Remarkably, GC-rich genes exhibit several-fold to over 100-fold higher expression efficiency compared to GC-poor counterparts [86].

## 5.0 Conclusion

Several conclusions can be inferred from the results of this study. The selected MHC class I and MHC class II epitopes demonstrated antigenicity, were non-allergenic, and non-toxigenic, offering a 94.91% population coverage when used in multi-epitope vaccine (MEV) construction. Notably, the epitopes “LAPDRRVEI” and “IEYAITPEI” were identified as IFN-γ inducers, and all chosen epitopes for the MEV exhibited 100% conservation. The MEV protein construct consisted of 16.67% alpha-helices, 8.87% extended beta strands, 4.55% beta turns, and 69.91% random coils, and was found to be soluble, thermostable, and basic with a molecular weight of 47.343 kDa. Further, molecular dynamics simulations of the docked MEV and TLR complexes suggested that these complexes were energetically feasible and possessed high binding affinity. The eigenvalues indicated a lower likelihood of deformation during immunological activity. The MEV also induced robust IgM and IgG responses, alongside their isotypes, and activated CD4+ and CD8+ T-cells, NK cells, dendritic cells, macrophages, and cytokines including IFN-γ, TGF-β, IL-2, IL-10, and IL-12. While these in silico results are promising, they warrant further validation through in vitro, pre-clinical, and clinical studies.

## Acknowledgments

The authors are thankful to the staff of the Bacterial Vaccine Production Department, National Veterinary Research Institute Vom, Nigeria for their technical support during this research.

## Author contributions

**Conceptualization:** Elayoni E. Igomu, Paul H. Mamman

**Data curation:** Elayoni E. Igomu, David O. Ehizibolo

**Investigation:** Elayoni E. Igomu, Paul H. Mamman, Jibril Adamu

**Methodology:** Elayoni E. Igomu, David O. Ehizibolo, Paul H. Mamman, Jibril Adamu, Abubakar O. Woziri

**Resources:** Maryam Muhammad

**Software:** Elayoni E. Igomu, Abubakar O. Woziri

**Supervision:** David O. Ehizibolo, Paul H. Mamman, Jibril Adamu

**Writing – Review and editing:** Elayoni E. Igomu, John A. Benshak, David O. Ehizibolo, Paul H. Mamman, Maryam Muhammad, Manasa Y. Sugun, Rhoda Sam-Gyang

